# Bivalent and Broad Chromatin Domains Regulate Pro-metastatic Drivers in Melanoma

**DOI:** 10.1101/721480

**Authors:** Christopher Terranova, Ming Tang, Mayinuer Maitituoheti, Ayush T. Raman, Jonathan Schulz, Samir B. Amin, Elias Orouji, Katarzyna Tomczak, Sharmistha Sarkar, Junna Oba, Caitlin Creasy, Chang-Jiun Wu, Dongyu Zhao, Kaifu Chen, Lauren E. Haydu, Wei-Lien Wang, Alexander J. Lazar, Scott E. Woodman, Chantale Bernatchez, Kunal Rai

## Abstract

Chromatin deregulation is an emerging hallmark of cancer. However, the extent of epigenetic aberrations during tumorigenesis and their relationship with genetic aberrations are poorly understood. Using ChIP-sequencing for enhancers (H3K27ac and H3K4me1), promoters (H3K4me3), active transcription (H3K79me2) and polycomb (H3K27me3) or heterochromatin (H3K9me3) repression we generated chromatin state profiles in metastatic melanoma using 46 tumor samples and cell lines. We identified a strong association of NRAS, but not BRAF mutations, with bivalent states harboring H3K4me3 and H3K27me3 marks. Importantly, the loss and gain of bivalent states occurred on important pro-metastasis regulators including master transcription factor drivers of mesenchymal phenotype including *ZEB1, TWIST1, SNAI1* and *CDH1*. Unexpectedly, a subset of these and additional pro-metastatic drivers (e.g. *POU3F2, SOX9* and *PDGFRA)* as well as melanocyte-specific master regulators (e.g. *MITF, ZEB2*, and *TFAP2A)* were regulated by exceptionally wide H3K4me3 domains that can span tens of thousands of kilobases suggesting roles of this new epigenetic element in melanoma metastasis. Overall, we find that BRAF, NRAS and WT melanomas may use bivalent states and broad H3K4me3 domains in a specific manner to regulate pro-metastatic drivers. We propose that specific epigenetic traits – such as bivalent and broad domains – get assimilated in the epigenome of pro-metastatic clones to drive evolution of cancer cells to metastasis.

## Introduction

Melanoma is a deadly disease with an estimated 96,000 new cases each year^1–3^. While targeted therapy and immunotherapy have become the standard of care with significant improvement in clinical response, more than 7,000 patients still succumb to this disease per year due to primary or acquired resistance^4,5^. Therefore, it is critical to gain a deeper understanding of the disease biology to design more effective therapies.

Large scale efforts from consortiums such as The Cancer Genome Atlas (TCGA) have provided deeper understanding of molecular aberrations in the disease^6–8^. These studies identified critical somatic mutations in this disease that likely occur due to UV exposure. Among these, somatic mutations in important *bona fide* oncogenes and tumor suppressors, such as BRAF, NRAS, NF1, INK/ARF, PTEN and TP53, have been demonstrated to be well-chronicled drivers of this malignancy^6–8^. One of the important findings from these studies were genetic aberrations in several key epigenetic regulators such as EZH2, IDH1/2, ARID2, KMT2C and KMT2D^6–8^. Many of these proteins are enzymes that regulate covalent modifications of histones^7,9–13^. Although recent studies provide insight into the correlation of isolated histone marks, there are a myriad of possible combinatorial patterns of histone modifications, and it is these combinatorial states - not individual modifications - that dictate epigenetic status of associated genomic loci^14–16^. These observations suggest that epigenetic alterations, including those in histone modifications, may play important roles in melanomagenesis. Indeed, specific functional roles have been assigned to some of these players as well as histone variants such as macroH2A and H2A.z in melanomas^17,18^. In addition, our previous study showed alterations in specific chromatin states during transition from pre-malignant to malignant phenotype in melanoma^19^.

These studies provide strong rationale for a systematic mapping of epigenome to obtain comprehensive understanding of epigenetic elements that may act as driver events in specific melanoma tumors. This concept has been epitomized by the DNA methylation profiles in large number of tumors by the TCGA study which has provided key concepts about roles of this important epigenomic mark in cancer progression^11^. For example, a subset of cancer types are demonstrated to harbor hypermethylation phenotype, termed CpG Island Methylation Phenotype, which associated with mutations in IDH1/2 genes^11^. Chromatin state mapping in large number of tumors has the potential to identify similar concepts^20^. Furthermore, several projects such as ENCODE and Roadmap Epigenomics have cataloged extensive histone modification in normal human tissues and cell lines which allows for identification of cancer specific alternations in chromatin states^21,22^. Since epigenetic aberrations are reversible by targeting of their enzyme regulators, the chromatin mapping efforts are likely to identify potentially novel therapeutic strategy in specific genetic context. For example, our previous study suggested that HDAC inhibitors could be a good strategy to block pre-malignant to malignant transition in melanoma^19^.

In this study, we present a comprehensive chromatin state analysis of metastatic melanoma in 46 tumor samples (profiled by TCGA) and cell lines (Profiled by Cancer Cell Line Encyclopedia (CCLE) or internal efforts at MD Anderson) by performing ChIP-sequencing of 6 histone modification marks. Overall, investigation revealed a mechanism in which alterations of distinct chromatin states, including bivalent and exceptionally wide H3K4me3 and H3K27me3 domains, regulate key drivers of a mesenchymal/invasive phenotype, a network of genes that includes master transcription factors and melanocyte-specific regulators. Together, this study encompasses the most complete description regarding the epigenetic circuitry governing melanoma metastasis that can serve as an important resource for advancing the understanding of the melanoma epigenome.

## Results

### Bivalent polycomb repressive chromatin domains distinguish metastatic melanoma tumors based on mutational subtypes

Chromatin state profiling remains a powerful tool for determining the regulatory status of annotated genes and identifying novel elements in non-coding genomic regions^3,4^. Using ChIP-sequencing for enhancers (H3K27ac and H3K4me1), promoters (H3K4me3), active transcription (H3K79me2) and polycomb (H3K27me3) or heterochromatin (H3K9me3) repression coupled with tissue-matched mutation, transcriptomic and methylation data, we describe the cis-regulatory landscape across 46 melanoma samples. These constituted 20 metastatic melanoma tumors (profiled by the TCGA study^6^), 10 patient-derived melanoma short term cultures (passage n < 10; profiled by internal effort at MD Anderson; manuscript in preparation) and 16 established melanoma lines profiled by the Cancer Cell Line Encyclopedia/Sanger (CCLE)^23^ (Supplementary Table **1**). Using our primary cohort of 20 metastatic melanoma tumor samples we computed multiple chromatin state models (8-states through 30-states) with the ChromHMM algorithm (Fig**. 1a** and Supplementary Fig. **1a**). We chose an 18-state model because it is large enough to identify important functional elements while still being small enough to interpret easily. This model includes active (E1) and transcribed promoters (E2), harboring high levels of H3K4me3, H3K27ac, H3K4me1 without (E1) and with H3K79me2 (E2) within the TSS or TSS flanking regions (Supplementary Fig. **1b**); transcribed genes (E3 and E4); genic (E5, E6) and active enhancers (E7, E8) harboring high levels of H3K27ac and H3K4me1 with concomitant enrichment of H3K79me2 within (E5-E6) or outside (E7-E8) the TSS flanking regions; and heterochromatic (E10) or polycomb (E14) based repression harboring high levels of either H3K9me3 (E10) or H3K27me3 (E14) respectively. In addition, we also observed two prominent bivalent/poised states: first, harboring both H3K4me3 and high levels of H3K9me3 (E12, annotated as “bivalent H3K9me3”), and second, H3K4me3 and high levels of H3K27me3 (E13, annotated as “bivalent H3K27me3”). Overall, the chromatin state profiles in metastatic melanoma tumors are associated with both gene expression patterns and DNA methylation levels (Fig. **1b, c**). As expected, active promoters (E1 and E2) are associated with high levels of gene expression and low levels of methylation, whereas repressed states (E12 and E13) are associated with low levels of gene expression and high levels of methylation (Fig. **1b, c**). Consistent with the previous Roadmap epigenome analysis^21,22^ in normal samples, DNA methylation patterns also showed some dynamic patterns. For example, hypermethylation in active chromatin states E3 through E8 are also associated with high levels of transcriptional activation (Fig. **1b, c**).

This TCGA tumor cohort is inclusive of multiple melanoma subgroups^6^, including mutation subtypes (BRAF, NRAS, WT), transcriptomic subtypes (Immune, Keratin, MITF-low) and DNA-methylation subtypes (CpG, Hypermethlyated, Hypomethylated, Normal). Before comparing chromatin state data and molecular subtypes, we first ensured the generated ChIP-seq profiles could be uniquely mapped to the specific donor through CNV analysis. In 18 of 20 tumors, the CNV analysis from ChIP-seq data correlated with the CNV classification from TCGA (Supplementary Fig. **1c, d**). Projection of chromatin state data using Multidimensional Scaling (MDS) analysis revealed chromatin state E13 (bivalent state containing high levels of H3K27me3 and H3K4me3 and modest enrichment of H3K9me3) was able to separate NRAS-mutant melanoma tumors from BRAF-mutants and WT samples in the first dimension (Fig. **1d, e** and Supplementary Fig. **1e**). Moreover, differential analysis of bivalent chromatin states (E12 and E13) between mutational subtypes (BRAF vs NRAS vs WT) demonstrated NRAS-enrichment specific to bivalent H3K27me3 high (Fig. **1f-h** and Supplementary Fig. **2a-c**). Together, this data suggested that a substantial number of genomic loci in melanoma tumors harbor bivalent chromatin states. To determine whether this could be a reflection of tumor heterogeneity, we assessed the presence of these bivalent chromatin states in 10 melanoma short-term cultures (MSTC) and 16 commercially available cell lines from CCLE (Supplemental Table **1**), allowing us to eliminate the possibility of signals emerging from other cell types in the tumor microenvironment. Using Model Based Analysis of ChIP-seq (MACS), we identified all potential bivalent combinations by directly overlapping H3K4me3 peaks with either H3K27me3 peaks (bivalent H3K27me3), H3K9me3 peaks (bivalent H3K9me3) or H3K27me3 peaks + H3K9me3 peaks (bivalent H3K4/H3K9/H3K27me3) in each individual sample. Bivalent loci were further identified as “common” in tumors and cell lines if they were present in ∼50% of the samples in each subgroup (BRAF = 7/13, NRAS = 2/4, WT = 2/3, MSTC = 5/10 and CCLE = 8/16). In accordance with our chromatin state analysis, NRAS-mutants displayed the greatest number of bivalent H3K27me3 loci out of all the subgroups whereas WT samples displayed the least number of bivalent loci (Fig. **1i** and Supplementary Fig. **2d**). Importantly, both the MSTC and CCLE subgroups also displayed a large number of bivalent H3K27me3 loci that were shared with melanoma tumors (Fig. **1i, j** and Supplementary Fig. **2e-g**), further suggesting these domains are enriched in cancer cells and are not a product of tumor heterogeneity.

**Figure 1:**
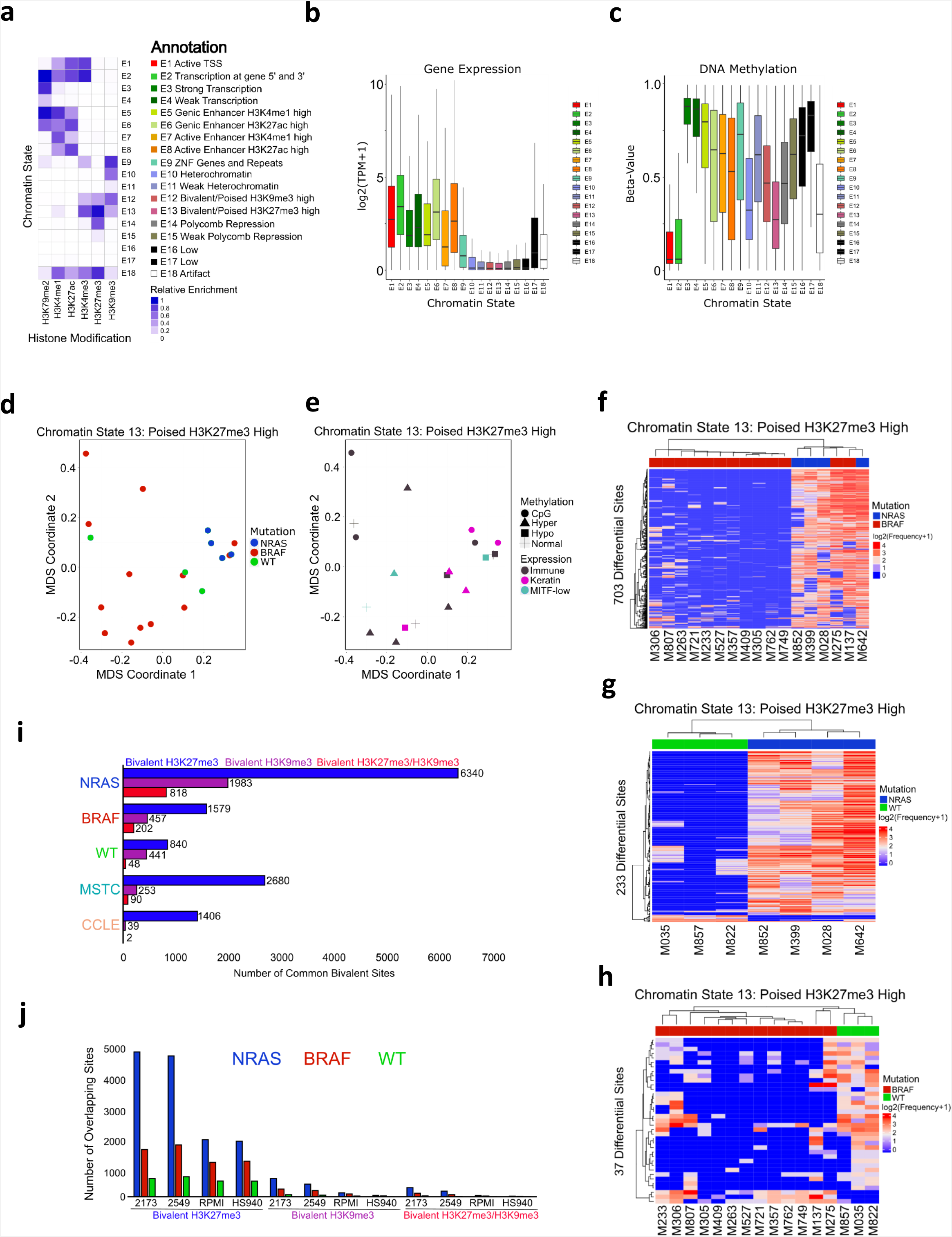
Bivalent chromatin domains distinguish metastatic melanoma tumors based on mutational subtypes. a) Combinatorial chromatin state definitions and histone mark probabilities identified in 20 metastatic melanoma tumor samples using the ChromHMM algorithm. b) Boxplot illustrating mean expression of genes from RNA-seq based on genomic regions overlapping with each chromatin state. c) Boxplot illustrating mean DNA methylation levels based on genomic regions overlapping with each chromatin state. d) MDS analysis of chromatin state E13 (poised H3K27me3 high) annotated by mutation (NRAS, BRAF, WT), (e) RNA-expression (Immune, Keratin, MITF-low) and DNA methylation (Normal, CpG, Hyper, Hypo) classifications from The Cancer Genome Atlas. f) Heatmap displaying differentially regulated regions (FDR < 0.05) of chromatin state E13 (poised H3K27me3 high) between NRAS and BRAF, (g) NRAS and WT and (h) BRAF and WT tumor subtypes. i) Common bivalent H3K27me3 (H3K4me3+H3K27me3), bivalent H3K9me3 (H3K4me3+H3K9me3) and bivalent H3K27/H3K9me3 (H3K4me3+H3K27me3+H3K9me3) binding sites in melanoma tumor subtypes, MSTC and CCLE lines. Binding sites were identified as common if they were present in at ∼50% of the samples (NRAS = 2/4, BRAF = 7/13, WT = 2/3, MSTC = 5/10 and CCLE = 8/16). j) Co-occupancy analysis of common bivalent binding sites in melanoma tumor subtypes directly overlapping bivalent binding sites in representative MSTC or CCLE lines.

**Figure 2:**
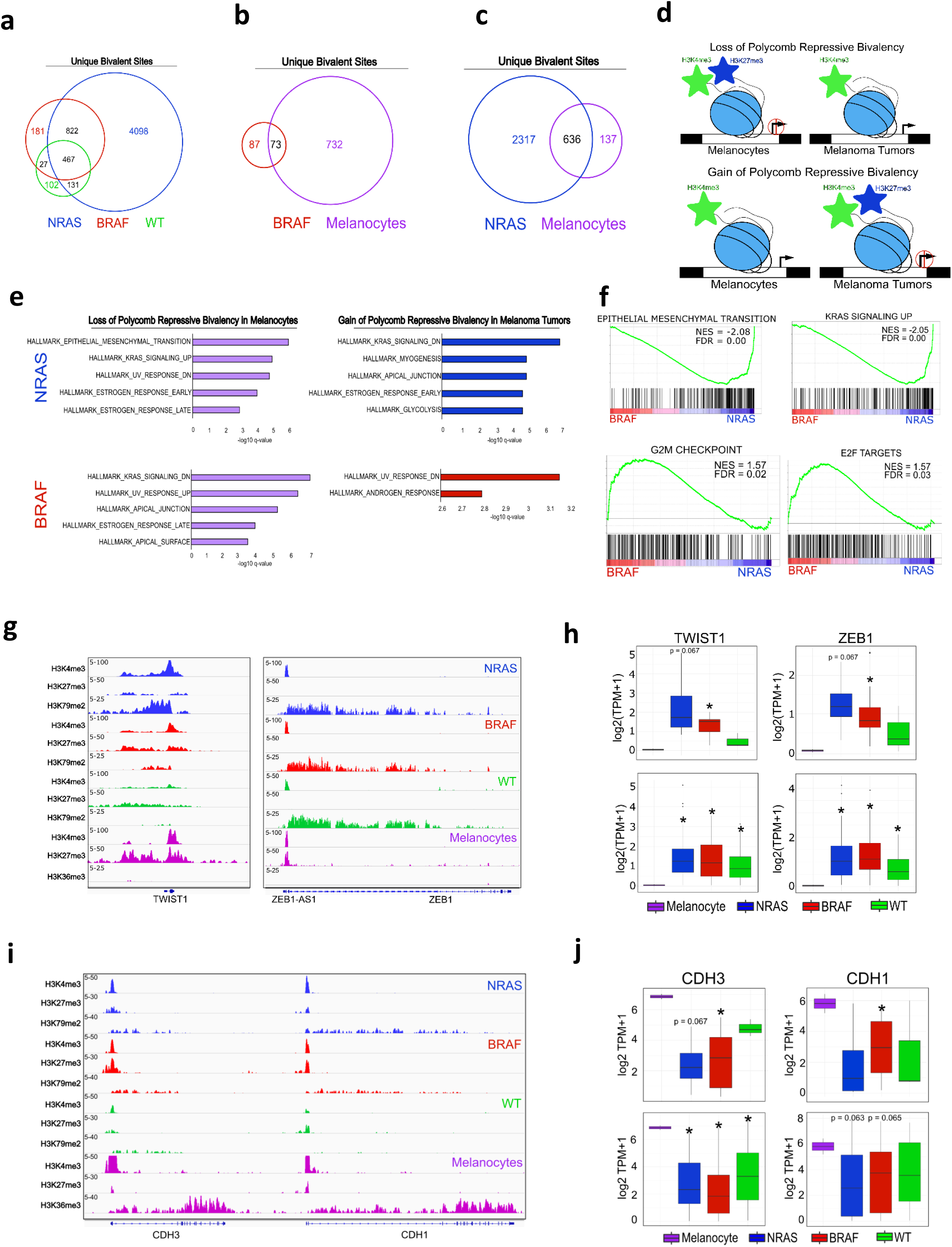
Bivalent domains are lost and gained on key mesenchymal genes in metastatic melanoma. a) Venn diagram analysis of bivalent H3K27me3 sites in NRAS, BRAF and WT tumor subtypes. b, c) Venn diagram analysis of unique bivalent H3K27me3 sites in NRAS and BRAF tumor subtypes overlapping bivalent polycomb sites in primary melanocytes from Roadmap. d) Schematic of bivalent H3K27me3 losses and gains in melanocytes vs melanoma tumors. e) Top 5 significant MSigDB/GSEA HALLMARK pathways based on bivalent H3K27me3 sites that are lost and gained within -/+10kbTSS-TES of a gene in NRAS and BRAF tumor subtypes. f) Significant GSEA HALLMARK pathways based on differentially expressed genes between NRAS (n = 81) vs BRAF (n = 118) metastatic melanoma tumor samples from TCGA. g) Genome browser view of ChIP-seq tracks for H3K4me3, H3K27me3 and active transcription (H3K79me2/H3K36me3) on the *TWIST1* and *ZEB1* genes in melanocytes and melanoma tumor subtypes. h) Boxplot displaying quantile normalized mean RNA-expression profiles (log2 TPM) of the *TWIST1* and *ZEB1* genes in melanocytes (n=2) and melanoma tumor subtypes (NRAS=4, BRAF=13, WT=3 (top), NRAS=81, BRAF=118, WT=38 (bottom). Asterisk denotes p-value < 0.05. i) Genome browser view of ChIP-seq tracks for H3K4me3, H3K27me3 and active transcription (H3K79me2/H3K36me3) on the *CDH3* and *CDH1* genes in melanocytes and melanoma tumor subtypes. j) Boxplot displaying quantile normalized mean RNA-expression profiles (log2 TPM) of the *CDH3* and *CDH1* genes in melanocytes (n=2) and melanoma tumor subtypes (NRAS=4, BRAF=13, WT=3 (top), NRAS=81, BRAF=118, WT=38 (bottom). Asterisk denotes p-value < 0.05.

### Bivalent domains are lost and gained on key mesenchymal genes in metastatic melanoma

In embryonic stem cells (ESCs), bivalent promoters mark critical lineage-specific genes which gain or lose these modifications as cells differentiate towards a particular phenotype^24,25^. Previous studies have demonstrated various cancer-related genes (i.e. CDKN2A) maintain or regain bivalent promoters in normal tissues, however their role in cancer progression has yet to be described^26^. To this end, we first computed unique bivalent loci (H3K4me3 and H3K27me3) in BRAF-mutant, NRAS-mutant and WT samples by overlapping common peaks in each tumor subtype which further suggested NRAS-mutants contained the highest number of bivalent loci (Fig. **2a** and Supplementary Fig. **3a-c**). To determine how subtype-specific bivalent polycomb losses and gains influence melanoma progression, we calculated the overlaps of bivalent sites from NRAS- and BRAF-mutants to those in primary melanocytes (Fig. **2b, c** and Supplementary Fig. **3d, e**). We focused primarily on the comparisons between NRAS- and BRAF-mutants as most significant differences in bivalency were observed to occur between these subgroups (Fig. **1f-i**). We posited that removal of H3K27me3 mark from bivalent loci in melanocytes would lead to transcriptionally ‘active’ loci in melanoma tumors and such loci were termed under “bivalent losses” (Fig. **2d**). Similarly, gain of H3K27me3 mark on loci (bivalent in tumors) that harbor only H3K4me3 in melanocytes will lead to transcriptional repression which were termed as “bivalent gains” (Fig. **2d**). Determination of the gene targets (within +/-10KB of each locus) and subsequent pathway enrichment analyses by GSEA MSigDB tool identified critical melanoma-associated “hallmark pathways” within each genetic subgroup (Fig. **2e**). For example, in NRAS-mutants, losses of melanocyte-specific bivalency included genes associated with the “epithelial-mesenchymal transition” and “KRAS-signaling up”, while gains of tumor-specific bivalency included genes associated with “KRAS-signaling down” and “apical junction” (Fig. **2e** and Supplementary Table **2**). Furthermore, RNA-seq analysis between NRAS-mutant (n = 81) and BRAF-mutant (n = 118) TCGA SKCM tumors confirmed “epithelial-mesenchymal transition” and “KRAS-signaling up” signatures to be upregulated specifically in NRAS-mutant samples (Fig. **2f** and Supplementary Table **3**), demonstrating an association between shifts in bivalent domains and gene expression patterns within critical melanoma pathways. Indeed, apart from obvious activation of RAS pathway genes, previous studies have demonstrated importance of activation of mesenchymal drivers and phenotypes in invasive behavior of metastatic melanoma and other malignancies^27–29^.

In order to identify important metastasis driver genes that are subjected to epigenetic regulation, we focused on genes that do not harbor genetic changes in cancers, which included key EMT transcription factors (EMT-TF) *ZEB1, TWIST1, SNAI1* and *CDH1* (Fig. **2e** and Supplementary Fig. **3f**). In response to activation of NRAS or BRAF, the EMT-TF network undergoes a reorganization that includes activation of *ZEB1* and *TWIST1* along with the loss of *CDH1*. This phenomenon is accompanied with increased invasion and correlates with poor prognosis in metastatic melanoma patients^28^. We observed that *ZEB1, TWIST1*, and additionally *SNAI1* and *TGFBI* are held in a bivalent state in melanocytes and transition to an active state in metastatic melanoma tumors, which occurs in a subtype specific manner (Fig. **2g** and Supplementary Fig. **3g**). Importantly, these events are significantly correlated with their mRNA expression levels (Fig. **2h** and Supplementary Fig**. 3h**). In addition, genes such and *CDH1* and *CDH3* harbor H3K4me3 in melanocytes and gain repressive bivalency in both NRAS-mutant and BRAF-mutant tumors subtypes (Fig. **2i**). This transition to repressive bivalency is associated with their downregulation (Fig. **2j**). Interestingly, *ZEB1* and *TWIST1* harbored bivalent chromatin states in embryonic stem cells and germ-layer stem cells, but not in mesenchymal stem cells where they are active (Supplementary Fig. **4a-d**). Various other tissues such as breast (*TWIST1* and *ZEB1*^*30,31*^), brain (*TWIST1* and *ZEB1*), colon (*TWIST1*), lung (*TWIST1*) and ovarian (*TWIST1*) normal tissues or cell lines (Supplementary Fig. **4a-d**) showed varying degree of bivalency which correlated well with gene expression patterns demonstrating this “bivalent loss” mechanism can be expanded to various other tissues. In addition to EMT-TF, we observed bivalent shifts on several well-characterized drivers of melanoma metastasis, such as *BMI1*^*32*^, *RNF2*^*33*^, and *CDK6*^*17*^ (Supplementary Fig. **3i-k**), further strengthening the hypothesis that shifts in bivalent domain can be a key epigenetic event during metastasis.

### Melanocyte-specific bivalent genes transition to transcriptionally active broad H3K4me3 in melanoma tumors

While surveying the transitions of bivalent domains to a transcriptionally active state through the genome, we observed two distinct types of H3K4me3 profiles present in many of the samples. For example, the *TWIST1* domain spanned well beyond the gene body whereas the *ZEB1* domain was localized within the promoter region (Fig. **2g** and Supplemental Fig. **4a, b**). In contrast to typical H3K4me3 domains that are usually 200-1000bp long, broad H3K4me3 domains can span thousands of kilobases and have been implicated in various cellular processes, including increased gene expression, enhancer activity and tumor-suppressor gene regulation^34–36^. Hence, we posited that some important driver genes that lose bivalency in melanocytes may retain or gain different types (broad or non-broad) of H3K4me3 domains in melanoma tumors.

Hence, we systematically identified broad H3K4me3 domains by computing the overall width and density from MACS2 broad peaks in BRAF-mutant, NRAS-mutant and WT tumors (Fig. **3a-c**). Globally, NRAS-mutant and BRAF-mutant subtypes harbored the largest number of broad H3K4me3 peaks, in some cases spanning >30kb in NRAS-tumors (Fig. **3b**). Similarly, the top 1% broadest domains in these subtypes extended well beyond the typical H3K4me3 domain (200-1000bp), with peaks reaching over 4kb in 10/17 the individual samples profiled (Fig. **3d**). In contrast, WT samples harbored much shorter H3K4me3 domains with the widest peaks spanning around 15kb (Fig. **3c**). Here, the top 1% broadest peaks did not markedly extend beyond the typical H3K4me3 domain, with only 1/3 samples harboring peaks over 4kb at this percentage cutoff (Fig. **3d**). Based on these observations and previous reports, broad domains were defined as peaks that extended at least 4x (≥ 4kb; merge size = 1kb) beyond that of typical H3K4me3 domain (Fig. **3d** and Supplementary Fig. **5a**). Using this method, we identified two distinct types of H3K4me3 in melanoma tumors including broad domains (≥ 4kb) spanning outside of the TSS (mean length = 12.3kb) and non-broad domains (< 4kb) localized within the promoter region (mean length = 2.2kb) (Fig. **3e-f** and Supplementary Fig. **5b**). GSEA analysis suggested occurrence of broad domains on genes regulating important biological processes implicated in melanoma metastasis and tumorigenesis, such as “TNFA signaling via NFKB” ^37^, “WNT signaling” ^38,39^, and Hedgehog signaling” ^40–42^ (Fig. **3g-i** and Supplemental Table **4**), further suggesting a role for broad domains in melanoma metastasis. Additionally, important melanoma drivers such as *NEAT1, MALAT1, MYC* and *EVX1* harbored broad H3K4me3 domains in and around their gene bodies (Fig. **3j**).

**Figure 3:**
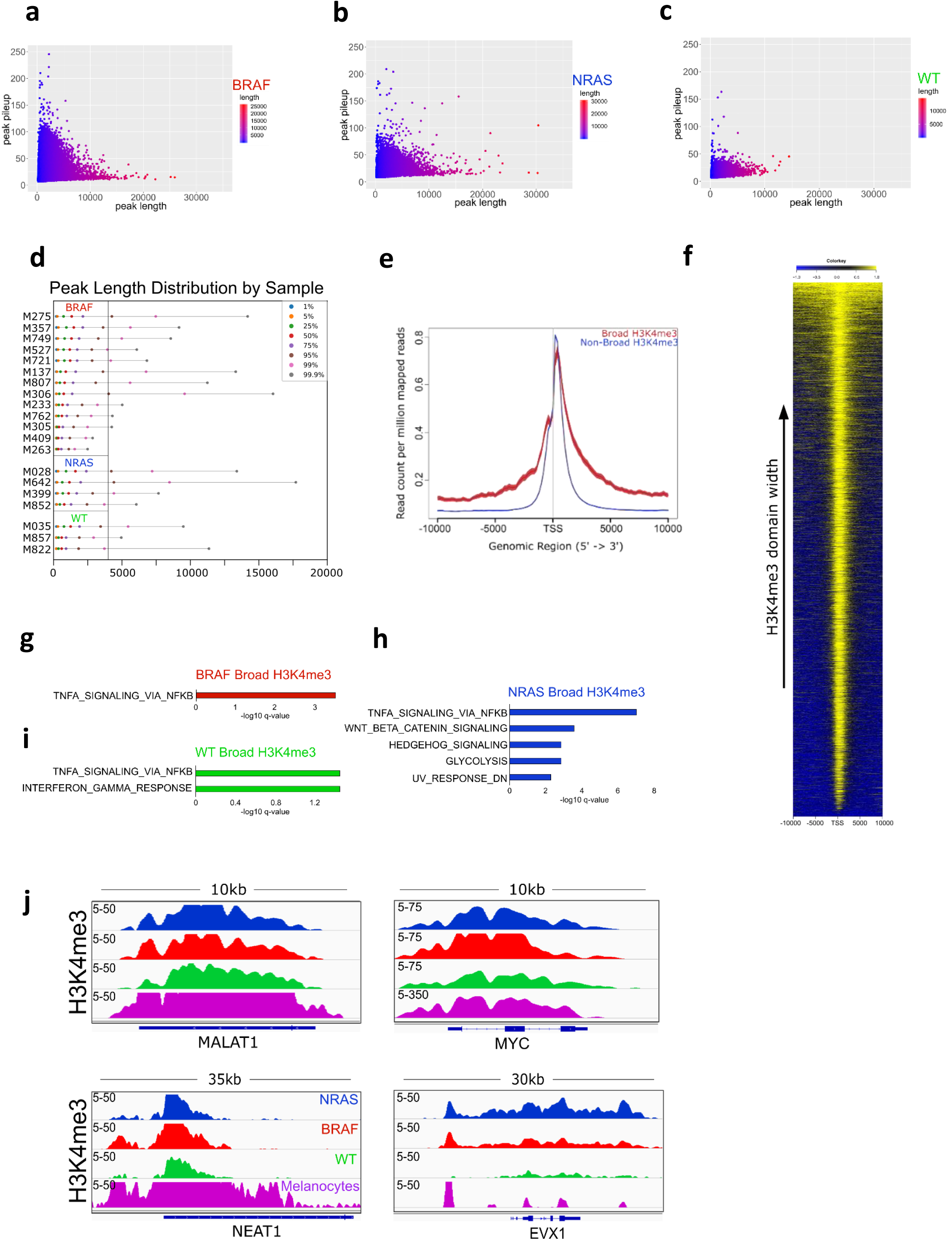
Distribution of exceptionally wide broad H3K4me3 domains in metastatic melanoma. a) Scatterplots of peak width (x-axis) and height (y-axis) from MACS2 broad peak calls (p-value 1e-5) for H3K4me3 in BRAF mutant tumors. b) Scatterplots of peak width (x-axis) and height (y-axis) from MACS2 broad peak calls (p-value 1e-5) for H3K4me3 in NRAS mutant tumors. c) Scatterplots of peak width (x-axis) and height (y-axis) from MACS2 broad peak calls (p-value 1e-5) for H3K4me3 in WT tumors. d) Quartile plot displaying peak length distribution based on percentage of broadest H3K4me3 domains in melanoma tumors. Black line denotes 4kb peak length. e) Average density profile displaying broad (>4kb) and non-broad (<4kb) H3K4me3 domains at 10kb to +10kb around TSS. f) Heatmap of H3K4me3 signal sorted by width at −10kb to +10kb around transcription start sites (TSS). g) Top significant MSigDB/GSEA HALLMARK pathways based on broad H3K4me3 domains within -/+10kbTSS of a gene in BRAF-mutant tumors. h) Top significant MSigDB/GSEA HALLMARK pathways based on broad H3K4me3 domains within -/+10kbTSS of a gene in NRAS-mutant tumors. i) Top significant MSigDB/GSEA HALLMARK pathways based on broad H3K4me3 domains within -/+10kbTSS of a gene in WT tumors. j) Genome browser view of ChIP-seq tracks for H3K4me3 on the *MALAT1, NEAT1, MYC* and *EVX1* genes in melanocytes and melanoma mutational subtypes.

To identify the subsets of melanocyte-specific bivalent genes that transition to transcriptionally active broad domains (and lose H3K27me3 in tumors), we further overlapped genes that harbor 1) bivalent domains uniquely in melanocytes (but not in tumors), 2) tumor-specific broad or non-broad H3K4me3 domains, 3) active transcription mark H3K79me2 and 4) gene expression [using RNA-seq data from NRAS (n=81), BRAF (n=118) and WT (n=38) metastatic samples]. Integrative analysis revealed NRAS-specific broad domains were associated with increased gene expression (Fig. **4a,b** and Supplemental Table **5**) and enriched for melanoma pathways such as “UV response up”, “KRAS signaling up” and “Glycolysis” (Fig. **4c**). Genes displaying the transition to transcriptionally active broad H3K3me3 included additional metastatic drivers known to function in the switch to an mesenchymal/invasive state, including *SOX9, POU3F2, PDGFRA* and *MYCN* (3.7kb width) (Fig. **4d, f**), While we also identified a broad H3K4me3-associated increase of transcription in WT samples, this occurred on a markedly lower number of genes many of which were shared with NRAS samples (Fig. **4a, b**). In contrast to broad domains, we identified a relatively large and constant number of genes transitioning from a bivalent state to non-broad H3K4me3 in melanoma tumors, many of which were present within all mutational subtypes (Fig. **4a**). On a global level, we did not observe a change in the mean expression levels of melanocyte-specific bivalent genes that transitioned to non-broad H3K4me3 domains in any melanoma subtype (Fig. **4b**), however, gene expression changes in specific genes were noted. These included various genes known to be misregulated upon the aberrant activation of RAS/RAF signaling, including *VCAN* as well as various RAS-effectors such as *RASGEF1B, RASEF, RASGRP1, RAPGEF5*, and *ARHGAP27* (Fig. **4e, g** and Supplementary Fig. **5c-e**), Overall, these results suggest that melanocyte-specific bivalent losses that transition to broad H3K4me3 domains in NRAS-mutant tumors regulate/increase the expression of mesenchymal/invasive drivers during metastatic progression.

**Figure 4:**
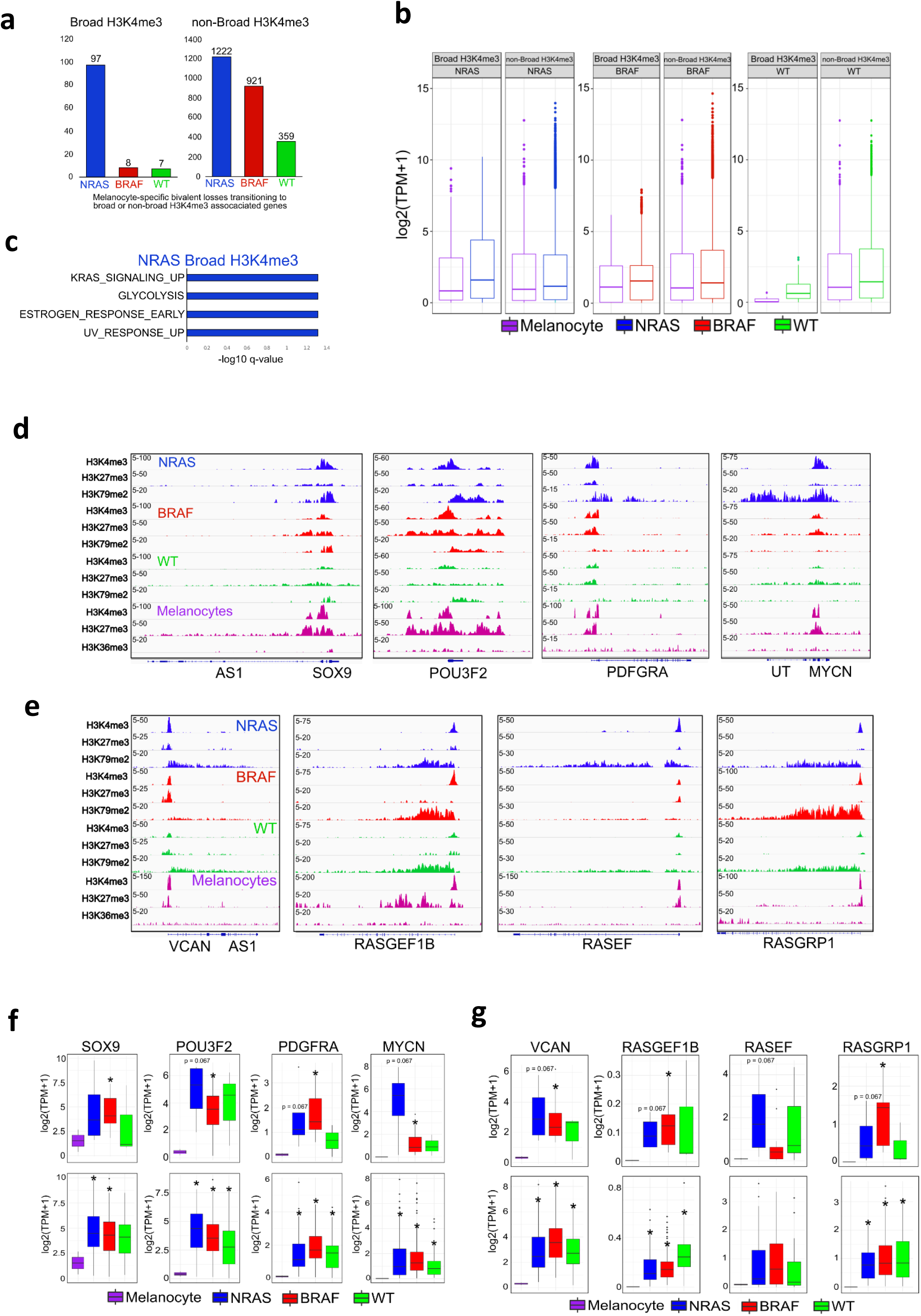
Melanocyte-specific bivalent genes transition to transcriptionally active broad H3K4me3 in NRAS mutant subtypes. a) Barplot of bivalent H3K27me3 genes (-/+10kbTSS-TES) that are lost in melanocytes and associated with broad or non-broad H3K4me3 in NRAS, BRAF and WT tumor subtypes. b) Barplot displaying quantile normalized mean RNA-expression profiles of NRAS (n=81), BRAF (n=118) and WT (n = 38) tumor subtypes based on broad or non-broad H3K4me3 genes identified in (a). c) Top significant GSEA HALLMARK pathways based on bivalent H3K27me3 genes (-/+10kbTSS-TES) that are lost in melanocytes and associated with broad H3K4me3 in NRAS tumor subtypes identified in (a) d) Genome browser views of ChIP-seq tracks for H3K4me3, H3K27me3 and active transcription (H3K79me2/H3K36me3) on the *SOX9, POU3F2, PDGFRA* and *MYCN* genes in melanocytes and melanoma tumor subtypes. e) Genome browser views of ChIP-seq tracks for H3K4me3, H3K27me3 and active transcription (H3K79me2/H3K36me3) on the *VCAN, RASGEF1B, RASEF* and *RASGRP1* genes in melanocytes and melanoma tumor subtypes. f) Boxplot displaying quantile normalized mean RNA-expression profiles (log2 TPM) of the *SOX9, POU3F2, PDGFRA* and *MYCN* genes in melanocytes (n=2) and melanoma tumor subtypes (NRAS=4, BRAF=13, WT=3 (top), NRAS=81, BRAF=118, WT=38 (bottom). Asterisk denotes p-value < 0.05. g) Boxplot displaying quantile normalized mean RNA-expression profiles (log2 TPM) of the *VCAN, RASGEF1B, RASEF* and *RASGRP1* genes in melanocytes (n=2) and melanoma tumor subtypes (NRAS=4, BRAF=13, WT=3 (top), NRAS=81, BRAF=118, WT=38 (bottom).

### Broad H3K4me3 domain spreading is associated with increased transcriptional activation

Our results suggest that during the transition from a bivalent to an active state, genes that retain broad H3K4me3 domains are associated with increased transcriptional activation compared to non-broad domains in melanoma tumors (Fig. **4a-c**). However, this was only observed on a small subset of genes (NRAS = 97; Supplementary Table **5**). We considered that another mechanism of gene activation could be spreading of the H3K4me3 signal, while gene repression may be associated with shortening of the broad H3K4me3 domains. Indeed, a previous report demonstrated that H3K4me3 domain shortening is associated with decreased gene expression of tumor suppressor genes in lung and liver cancers^36^. Therefore, we investigated whether the size of H3K4me3 domains are altered in metastatic melanoma. First, on a genome-wide level, we observed preferential shortening (≤ 2kb) of all H3K4me3 peaks in melanoma tumors relative to melanocytes (Fig. **5a**). Focusing on the promoter-associated broad H3K4me3 peaks (-/+10kb TSS), we further observed preferential shortening (≤ 2kb) of these peaks in melanoma tumors which was associated with a decrease in gene expression levels (Fig. **5b-d** and Supplemental Table **6**). Promoters harboring some of the broadest H3K4me3 domains displayed marked shortening on critical components of the melanocyte regulatory network in melanoma tumors, including *PMEL (aka GP100), MITF, ZEB2* and *TFAP2A*^*43*^ (Fig. **5e** and Supplementary Fig. **6a, b**). Because genes such as *MITF* and *ZEB2* are differentially expressed in alternate phenotypic states, displaying high expression in melanocytes (or proliferative phenotype) and low expression in melanoma tumors^28,44^ (Fig. **5f** and Supplementary Fig. **6c**), this finding illustrates a new epigenetic mechanism for their misregulation in metastatic melanoma. Similar to that of bivalent losses, we also identified a smaller subset of promoters that gained broad H3K4me3 in melanoma tumors. The top promoters displaying a broad H3K4me3 transition (≥ 10kb, 29 genes) were enriched for critical developmental regulators including *DLX1/2, TBX3, HMX2/3* and *LBX1*, many of which were also upregulated in metastatic melanoma tumors and hence suggest potential oncogenic roles for these proteins (Fig. **5g, h** and Supplementary Fig. **6d, e**). Together, these results suggest broad H3K4me3 domains are associated with higher expression that decreases upon shortening on key melanocyte-specific promoters during the transition to metastatic melanoma.

**Figure 5:**
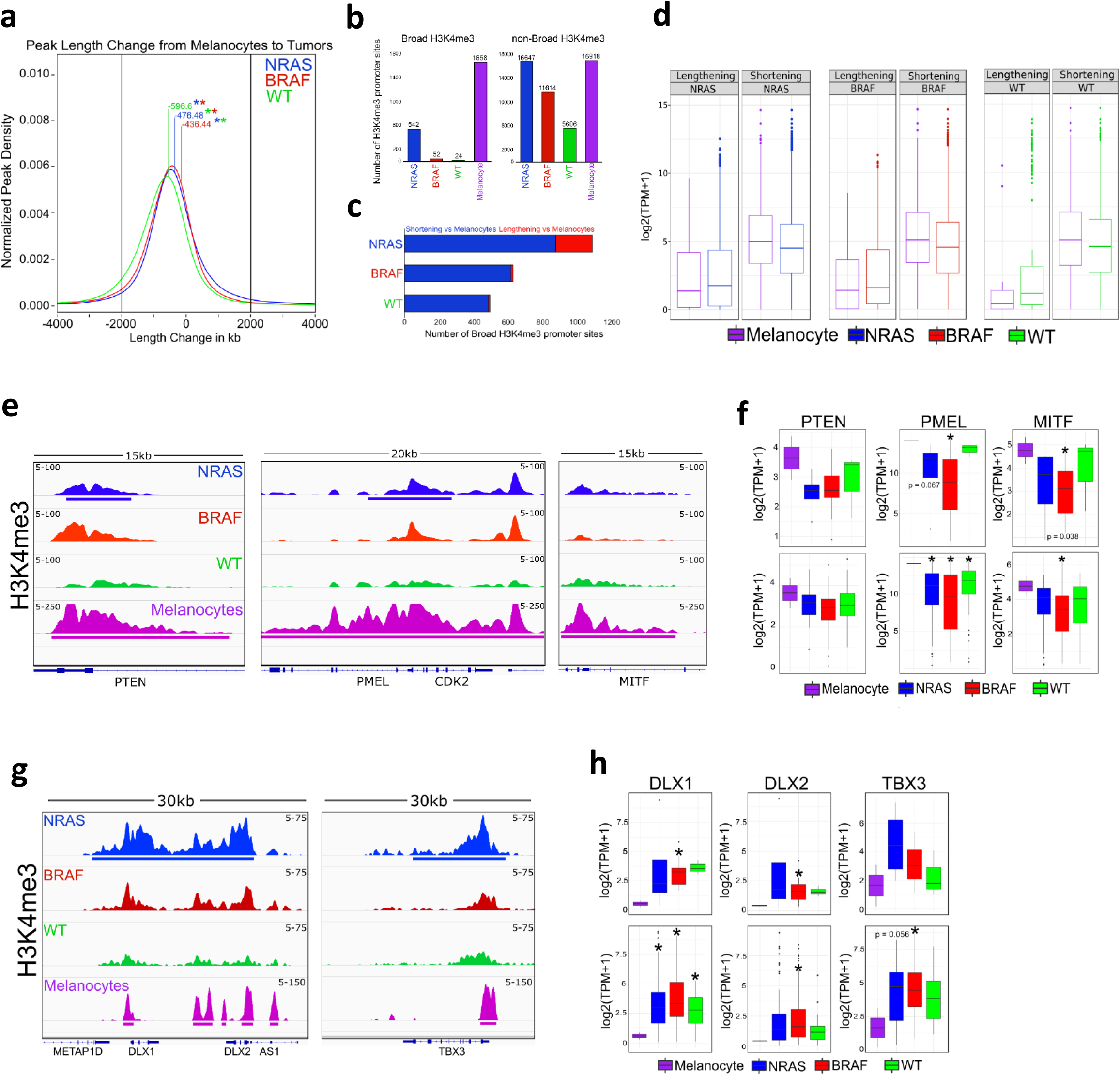
Broad H3K4me3 domain spreading is associated with increased transcriptional activation. a) Kernel density estimation plot displaying H3K4me3 peak length change (-/+2kb) from melanocytes to mutational subtype. Number denotes mean length change in kilobases between melanocytes and melanoma subtype. Asterisk denotes p-value < 1e-50 between melanoma subtype. b) Barplot of broad and non-broad H3K4me3 promoter associated sites in each mutational subtype. c) Barplot of broad H3K4me3 promoter associated sites displaying shortening or lengthening (-/+2kb) in melanoma tumors relative to melanocytes. d) Boxplot displaying quantile normalized mean RNA-seq expression profiles from NRAS (n=81), BRAF (n=118) and WT (n=38) metastatic samples based on genes displaying shortening or lengthening (-/+2kb) in melanoma tumors relative to melanocytes identified in (c). e) Genome browser view of ChIP-seq tracks displaying H3K4me3 shortening on the *PTEN, PMEL* and *MITF* genes in melanoma tumors relative to melanocytes. f) Boxplot displaying quantile normalized mean RNA-expression profiles (log2 TPM) of the *PTEN, PMEL* and *MITF* genes in melanocytes (n=2) and melanoma tumor subtypes (NRAS=4, BRAF=13, WT=3 (top), NRAS=81, BRAF=118, WT=38 (bottom). Asterisk denotes p-value < 0.05. g) Genome browser view of ChIP-seq tracks displaying H3K4me3 lengthening on *DLX1/2* and *TBX3* genes in melanoma tumors relative to melanocytes. h) Boxplot displaying quantile normalized mean RNA-expression profiles (log2 TPM) of the *DLX1/2* and *TBX3* genes in melanocytes (n=2) and melanoma tumor subtypes (NRAS=4, BRAF=13, WT=3 (top), NRAS=81, BRAF=118, WT=38 (bottom). Asterisk denotes p-value < 0.05.

### Broad H3K27me3 domains display preferential lengthening in metastatic melanoma

Another possible transition of bivalent domains is the loss of H3K4me3 and retention or extension of H3K27me3 domains. Therefore, we identified H3K27me3 domains by computing overall width and density from MACS2 broad peaks in BRAF-mutant, NRAS-mutant and WT samples (Supplementary Fig. **7a-c**). In all subtypes, we observed exceptionally wide H3K27me3 domains spanning tens of thousands of kilobases with multiple peaks extending well over 100kb in BRAF- and NRAS-mutant samples (Supplementary Fig. **7a-c**). Globally, the top 1% broadest H3K27me3 domains in 14/20 tumors extended well over 10kb (Fig. **6a** and Supplementary Fig. **7e**; merge size = 1kb). In contrast to the results for H3K4me3, we observed preferential lengthening of H3K27me3 domains on a global level (≥ 2kb) in BRAF-, NRAS-mutants and WT samples (Fig. **6b**). However, since H3K27me3 is already known as a broad mark, we focused on promoter regions that displayed large extensions or constrictions (≥ 10kb) of H3K27me3 domains. Although we observed relatively consistent number of broad (≥ 4kb) H3K27me3-associated promoters across mutational subtypes (Fig. **6c**), each subtype displayed a different pattern of H3K27me3 in comparison to melanocytes, with NRAS- and BRAF-mutant samples displaying preferential lengthening and WT preferential shortening (Fig. **6d**). Importantly, melanocytes harbored much shorter broad H3K27me3 domains (∼50kb) compared to all tumors (Supplemental Fig**. 7d**), While the largest constriction of H3K27me3 was just over 50kb, the largest extension spanned over 100kb in BRAF- and 85kb in NRAS-mutant samples (Supplemental Table **7**), further indicating polycomb domains are exceptionally broad in melanoma tumors. Overall, the widest H3K27me3 domain lengthening observed was on the *HOXC* locus spanning 102.8kb in BRAF samples (Fig. **6e**). Interestingly, as 3’ *HOX* genes are known to be expressed first during normal development, we observed H3K27me3 spreading specifically towards the 3’ end of the *HOXC* cluster in melanoma tumors. This 5’ -> 3’ H3K27me3 spreading was also observed on other *HOX* cluster genes (*HOXB*/*HOXD*) and further associated with the downregulation of various 3’ members relative to melanocytes (Fig. **6f-h** and Supplementary Fig. **7f, g**). Together, this exceptionally wide H3K27me3 spreading illustrates a new epigenetic signature for gene silencing of critical regulatory genes in metastatic melanoma.

**Figure 6:**
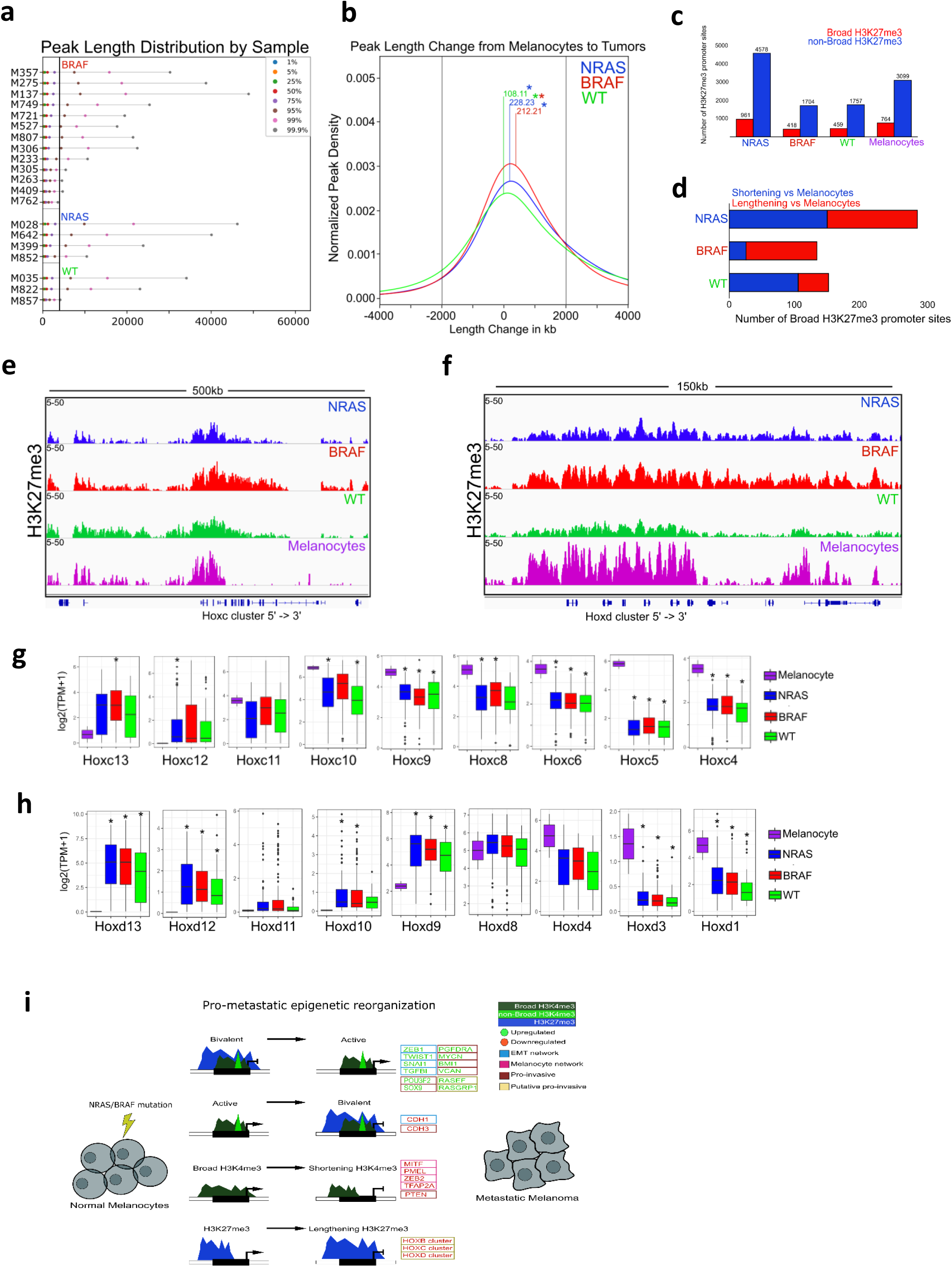
Broad H3K27me3 domains display preferential lengthening in metastatic melanoma. a) Quartile plot displaying peak length distribution based on percentage of broadest H3K27me3 domains in melanoma tumors. Black line denotes 4kb peak length. b) Kernel density estimation plot displaying H3K27me3 peak length change (-/+2kb) from melanocytes to mutational subtype. Number denotes mean length change in kilobases between melanocytes and melanoma subtype. Asterisk denotes p-value < 1e-15 between melanoma subtypec) Barplot of broad (>4kb) and non-broad (<4kb) H3K27me3 promoter associated sites in each mutational subtype. d) Barplot of broad H3K4me3 promoter associated sites displaying shortening or lengthening (>10kb) in melanoma tumors relative to melanocytes. e) Genome browser views of ChIP-seq tracks displaying exceptionally wide H3K27me3 lengthening from the 5’->3’ end on *HOXC* cluster genes in BRAF mutational subtypes. f) Genome browser views of ChIP-seq tracks displaying exceptionally wide H3K27me3 lengthening from the 5’->3’ end on *HOXB* cluster genes in NRAS mutational subtypes. g) Boxplot displaying quantile normalized mean RNA-expression profiles (log2 TPM) of all *HOXC* cluster genes in melanocytes (n=2) and melanoma tumor subtypes (NRAS=81, BRAF=118, WT=38). Asterisk denotes p-value < 0.05. h) Boxplot displaying quantile normalized mean RNA-expression profiles (log2 TPM) of all *HOXD* cluster genes in melanocytes (n=2) and melanoma tumor subtypes (NRAS=81, BRAF=118, WT=38). Asterisk denotes p-value < 0.05. i) Model for potential mechanism in which bivalent switches, the shortening of H3K4me3 domains and lengthening of H3K27me3 domains facilitate the reorganization of an interconnected regulatory network encompassing EMT-TF, melanocyte master regulators and mesenchymal/invasive genes to promote melanoma metastasis

## Discussion

Our study describes an epigenetic program for reorganization of an interconnected regulatory network encompassing EMT-TF, melanocyte master regulators and mesenchymal/invasive genes as melanocytes progress to metastatic melanoma. Our work provides three major advances regarding the role of the epigenome in melanoma metastasis: 1) Specific genetic events such as BRAF or NRAS mutations may utilize specific chromatin states to bring about transcriptional changes unique that genotype. 2) Chromatin state switches involving bivalent domains mark master regulatory genes associated with a metastatic phenotype, including the epithelial-to-mesenchymal transition. 3) In addition to bivalent domain switches, contraction or extension of broad H3K4me3 and H3K27me3 domains may be a key driver in the regulation of pro-metastatic genes. In addition to these findings, the current study encompasses the most comprehensive dataset from a large number of well-annotated samples which can help the community better understand the epigenetic circuitry governing melanoma metastasis.

Our results suggest that assimilation of epigenetic traits may be critical to evolution of metastatic clones, consistent with some prior studies^17,45^. It has been argued that evolution of cancer cells through metastasis entails constant acquisition and switching of cellular phenotypes during its journey from the primary site to colonization in a distant organ^46^. Hence, the molecular processes governing metastasis may mirror those driving tissue differentiation during normal development. At the same time, those molecular traits that allow establishment/evolution of neo-phenotypes, such as the ones in play during selection of *crossweinless* phenotype observed by Waddington many decades ago^47^, may also play an important driver role in evolution of cells undergoing metastasis. Waddington’s “genetic assimilation” model suggested that specific traits (genetic or epigenetic) driving a cell towards a specific differentiation state in response to certain environmental pressure are assimilated in the genome (or epigenome) during evolution. Similarly, we propose that during evolution of cancer cells to metastasis, under selective environmental pressure specific cell clones assimilate epigenetic traits – such as shifts in bivalent and broad domains – in their genome which eventually help them disseminate and colonize distant organs.

In ESCs, bivalent promoters mark critical lineage-specific genes which gain or lose these modifications as cells differentiate towards a particular phenotype^24,25^. Moreover, various cancer-related genes maintain or regain bivalent promoters in normal tissues^26^. Here, we expand on those studies by demonstrating that bivalent domains are lost and gained on critical EMT-TF, melanoma-specific drivers and mesenchymal genes as melanocytes progress to metastatic melanoma. Shifts from bivalent states to active or repressive marks are likely the least energy-intensive route to gene activation or repression respectively (https://doi.org/10.1101/456525). Since metastasis involves several cell fate transitions, which likely employ transcriptional circuitry consisting of multiple genes, energetically the use of shifts involving bivalent chromatin states would be favored as the preferred mode of gene regulation. It can be suggested that this specific epigenetic process could constitute one of the primary ‘forces’ that can help with the ‘canalization’ (per Waddington model) of the pro-metastatic clones.

Interestingly, our study for the first time suggests that melanomas harbor exceptionally wide H3K4me3 domains on key drivers of invasion and metastasis (*POU3F2*^48,43^, *SOX9*^49^, *PDGFRA*^50^ and *PDGFA)* and this transition was associated with increased transcriptional activation. It is likely, barring physical and chemical barrier, that addition of H3K4me3 marks on adjacent nucleosome may require less energy due to already recruited methylation machinery, and hence may be a preferred mode for relative increase in gene expression to meet the needs of metastatic cells. The observations of broad domains on critical components of the melanocyte regulatory network, including *MITF*^51,52^, *ZEB2*^28,44^, *TFAP2A*^43^ and *PMEL (aka GP100)*, as well as other genes not known to be previously associated with melanoma, including *MYCN, KLF6, NR2F2 and CPEB2,* may allow for identification of pro-metastatic drivers whose expression is mediated by epigenetic events. Overall, our work identifies new principles of epigenome regulation in melanoma metastasis and strengthens the hypothesis that metastatic dissemination is likely driven by specific epigenetic events.

## Supporting information

Supplemental_Table1_PatientInformation

Supplemental_Table2_BivalentGainsLosses_MSigDB_Pathways

Supplemental_Table3_TCGA_SKCM_BRAFvsRAS_RASlevel_DEGs

Supplemental_Table4_BroadH3K4me3_MSigDB_Pathways

Supplemental_Table5_BivalentLoss_to_BroadH3K4me3_MSigDB_Pathways

Supplemental_Table6_BroadH3K4me3_Shortening_Lengthening

Supplemental_Table7_BroadH3K27me3_Shortening_Lengthening

## Author contributions

CT and KR conceptualized and designed the study; CT and MM generated the ChIP-seq data; CT, MT, AR, JS and SA processed and performed data analysis for ChIP-seq; CT, MT and AR processed and performed data analysis for RNA-seq; JO, CC, CJW, CB and SW generated and characterized the Short Term Culture datasets; EO, SS, KT, DZ and KC provided intellectual input; CT and KR wrote and prepared the manuscript. All authors read and approved the manuscript.

## Acknowledgements

We are grateful to Kadir C. Akdemir, Yonathan Lissanu Deribe, Anand Singh, Scott Callahan, Veena Kochat, Sharon Landers, Angela Bhalla and Keila Torres for helpful discussions and proof reading the manuscript. The work described in this article was supported by grants from the National Institutes of Health (CA160578 to K. R.; CA157919 and CA016672 to SMF Core), Center for Cancer Epigenetics at MDACC (K.R.) and MD Anderson Cancer Center (Start-up funds to K.R.). STR DNA fingerprinting was done by the UTMDACC CCSG-funded Characterized Cell Line Core, NCI # CA016672. The molecular characterization of the short-term culture tumor lines was supported by generous philanthropic contributions to The University of Texas MD Anderson Moon Shots Program.

## Methods

### Collection of melanoma tumor samples

Melanoma tumors were obtained from the Melcore tumor bank at MD Anderson Cancer Center.

### Characterization of melanoma short term cultures

Melanoma short term cultures were generated from metastatic tumor specimens as part of the Adoptive T-cell Therapy Clinical Program at the University of Texas, MD Anderson Cancer Center (LAB06-0755 and 2004-0069), as previously described^53^,^54^,^55^). Briefly a tumor specimen from metastatic tumor was collected and incubated with an enzymatic digestion cocktail (0.375% collagenase type I, 75 µg/ml hyaluronidase and 250 U/mL DNAse I) in tumor digestion medium (RPMI, 10 mM of HEPES, 1x Penicillin-Streptomycin and 20 ug/mL of Gentamicin; Gibco/Invitrogen) in a humidified incubator at 37°C with 5% CO_2_ and with a gentle rotation for 2-3h to obtain a single cell suspension. The tumor digest was filtered through a 70-µm filter, washed, and re-suspended in a serum free media, which was then plated in one well of a 6-well culture plate and incubated at 37 °C. After 24h, the media was replaced with fresh tumor media, comprised of RPMI with GlutaMAX, 10% FBS, Penicillin/Streptomycin, Gentamicin, β-mercaptoethanol (50 uM, Gibco/Invitrogen), HEPES (10 mM), and insulin-selenium-transferin (5 ug/ml, Gibco/Invitrogen). Cells were grown in enriched DMEM/F12 culture media (Gibco/Invitrogen) supplemented with all growth factors including 10%FBS, sodium pyruvate (1mM), insulin-selenium-transferin, MycoZap-PR (Lonza), HEPES (10mM) and β-Mercaptoethanol. Once enough cells were grown, the purity of the tumor was tested using a melanoma tumor surface marker (MCSP-1) by flow cytometry. Cultures were deemed established when the cells stained positive for a melanoma tumor marker (MCSP-1) and negative for a fibroblast marker (CD90). Appropriate serum starving was performed to eliminate fibroblasts. All cell lines were tested for mycoplasma using MycoAlert detection kit (Lonza), and fingerprinted by STR fingerprinting, and validated by comparing with matched blood samples. A few passages after a pure tumor line was established, the cells were cryopreserved and kept in stocks in liquid nitrogen until use.

### Chromatin immunoprecipitation assays

ChIP assays were performed as described previously^56^. Briefly, ∼2 × 107 cells were harvested via cross-linking with 1% (wt/ vol) formaldehyde for 10 min at 37 °C with shaking. After quenching with 150 mM glycine for 10 min at 37 °C with shaking, cells were washed twice with ice-cold PBS and frozen at −80 °C for further processing. Cross-linked pellets were thawed and lysed on ice for 30 min in ChIP harvest buffer (12 mM Tris-Cl, 1 × PBS, 6 mM EDTA, 0.5% SDS) with protease inhibitors (Sigma). Lysed cells were sonicated with a Bioruptor (Diagenode) to obtain chromatin fragments (∼200–500 bp) and centrifuged at 15,000 × g for 15 min to obtain a soluble chromatin fraction. In parallel with cellular lysis and sonication, antibodies (5 μg/3 × 106 cells) were coupled with 30 μl of magnetic protein G beads in binding/blocking buffer (PBS + 0.1% Tween + 0.2% BSA) for 2 h at 4 °C with rotation. Soluble chromatin was diluted five times using ChIP dilution buffer (10 mM Tris-Cl, 140 mM NaCl, 0.1% DOC, 1% Triton X, 1 mM EDTA) with protease inhibitors and added to the antibody-coupled beads with rotation at 4 °C overnight. After washing, samples were treated with elution buffer (10 mM Tris-Cl, pH 8.0, 5 mM EDTA, 300 mM NaCl, 0.5% SDS), RNase A, and Proteinase K, and cross-links were reversed overnight at 37. Immune complexes were then washed five times with cold RIPA buffer (10mM Tris–HCl, pH 8.0, 1mM EDTA, pH 8.0, 140mM NaCl, 1% Triton X-100, 0.1% SDS, 0.1% DOC), twice with cold high-salt RIPA buffer (10mM Tris–HCl, pH 8.0, 1mM EDTA, pH 8.0, 500mM NaCl, 1% Triton X-100, 0.1% SDS, 0.1% DOC), and twice with cold LiCl buffer (10mM Tris–HCl, pH 8.0, 1mM EDTA, pH 8.0, 250mM LiCl, 0.5% NP-40, 0.5% DOC). ChIP DNA was purified using AMPure XP beads (Agencourt) and quantified using the Qubit 2000 (Invitrogen) and Bioanalyzer 1000 (Agilent). Libraries for Illumina sequencing were generated following the New England BioLabs (NEB) Next Ultra DNA Library Prep Kit protocol. A total of 10 cycles were used during PCR amplification for the generation of all ChIP-seq libraries. Amplified ChIP DNA was purified using double-sided AMPure XP to retain fragments (∼200–500 bp) and quantified using the Qubit 2000 and Bioanalyzer 1000 before multiplexing.

### ChIP-seq data processing

Raw fastq reads for all ChIP-seq experiments were processed using a snakemake based pipeline^57^. Briefly, raw reads were first processed using FastQC (http://www.bioinformatics.babraham.ac.uk/projects/fastqc/) and uniquely mapped reads were aligned to the hg19 reference genome using Bowtie version 1.1.2 ^58^. Duplicate reads were removed using SAMBLASTER ^59^ before compression to bam files. To directly compare ChIP-seq samples uniquely mapped reads for each mark were downsampled per condition to 18 million, sorted and indexed using samtools version 1.5 ^60^. To visualize ChIP-seq libraries on the IGV genome browser, we used deepTools version 2.4.0 ^61^ to generate bigWig files by scaling the bam files to reads per kilobase per million (RPKM). Super ChIP-seq tracks were generated by merging bam files from each cancer type, sorting and indexing using samtools and scaling to RPKM using deepTools.

### Chromatin state calls

ChromHMM ^62^ was used to identify combinatorial chromatin state patterns based on the histone modifications studied. Normalized bam files were converted to bed files and binarized at a 1000bp resolution using the BinarizeBed command. We specified that ChromHMM should learn a model based on 18 chromatin states. As we considered models between 8 and 30 chromatin states, we chose an 18-state model because it is large enough to identify important functional elements while still being small enough to interpret easily. To determine chromatin state differences between different groups we used a two-step process. First, using the segmentation calls from the ChromHMM output the entire genome is divided into non-overlapping windows of 10 Kb. We next count the number of times a chromatin state is observed in each of the 10 Kb windows and obtain a frequency matrix for each state in the ChromHMM model (E1-E18). In the second step, low variable genomic loci are removed from the frequency matrix and significant differences between two groups of samples types are calculated by using a nonparametric Mann Whitney Wilcoxon test with a P-value < 0.05 for each state separately.

### Correlation of copy number from ChIPseq and SNP array

SKCM TCGA copynumber data were downloaded by TCGAbiolinks^63^. Copy number analysis for ChIPseq was carried out using copywriteR^64^, which uses off-target reads for accurate copy number detection. The ChIP-seq input files were used, which represent low-pass whole genome sequencing data sets. Bin size of 200kb was used for analysis. The copy number of each gene was determined by overlapping the genes with the segmentation files from ChIPseq and SNP array, respectively. The Pearson correlation of the copy number of all genes among samples was calculated and plotted in heatmap by ComplexHeatmap.

### Correlation of RNA expression and chromatin state data

SKCM TCGA RNAseq data were downloaded by TCGAbiolinks^63^. TPM (transcript per million) value was calculated from raw counts by scaling to the gene length first and then the library size. For each gene in a sample, the transcription start site is overlapped with the chromatin state segmentation file to determine the state of that gene. The expression values (TPM) for all genes and all samples are then combined and spilt by categories (18) of chromatin states. A box plot is plotted for each chromatin state.

### Correlation of DNA methylation and chromatin state data

SKCM TCGA 450k DNA methylation data were downloaded by TCGAbiolinks^63^. For each sample and each chromatin state segmentation bin, the average of beta values from the DNA methylation data were calculated for each bin. The average values of all bins from all samples are combined and then split by categories (18) of chromatin states. A box plot is plotted for each chromatin state.

### Identification and visualization of ChIP-seq binding sites

We used Model-based analysis of ChIP-seq (MACS) version 1.4.2 ^65^ peak calling algorithm to identify H3K4me3 (p-value of 1e-5) and MACS version 2.1.0 to identify H3K27me3 (p-value of 1e-5) enrichment over “input” background. Bivalent sites were identified by overlapping H3K4me3 with H3K27me3 or H3K9me3 by a minimum of 1bp using intersectBed^66^. Bivalent polycomb-heterochromatin regions were identified by overlapping the H3K4me3+H3K27me3 output with H3K9me3 by a minimum of 1bp. Common sites (NRAS = 2/4, BRAF = 7/13, WT = 2/3, MSTC = 5/10, CCLE = 8/16 and Melanocytes = 2/2) were identified using bedops^67^ with the following command; cat *bed | sort-bed - | bedmap --count --echo --delim ‘\t’ - | uniq | awk ‘$1 >=x’ | cut-f2-> samplename_common.bed. Final peaksets used for downstream analysis were generated using mergeBed. Unique BRAF, NRAS, WT and melanocyte peaks for bivalent polycomb, bivalent heterochromatin and bivalent polycomb-heterochromain were identified using the concatenate, cluster and subtract tools from the Galaxy/Cistrome web based platform ^68^. To identify sites that were bivalent in melanocytes but active in melanoma tumors, and visa-versa, common H3K27me3 sites were subtracted from the opposing bivalent site in each comparison.

### Identification of broad H3K4me3 and H3K27me3 domains

Broad H3K4me3 domains were identified using MACS2.1.0 with the broad setting (p-value of 1e-5) followed by merging adjacent peaks within 1kb using mergeBed^66,69^. We determined the optimal distance to merge adjacent peaks based on the number of broad H3K4me3 and H3K27me3 domains at distance thresholds between 1kb through 10kb in each mutational subtype. Broad H3K4me3 domains were further classified within the top 1% of the broadest peaks that extended at least 4x (≥4kb) beyond that of a typical H3K4me3 domain (0.2kb-1kb). Final peaksets for common sites broad and non-broad H3K4me3 and H3K27me3 domains were defined as peaks present in ∼50% of each mutational subtype and melanocytes (NRAS = 2/4, BRAF = 7/13, WT = 2/3 and Melanocytes = 2/2) as described in identification and visualization of ChIP-seq binding sites.

### Assigning binding sites to genes

A list of Ensembl genes was obtained from the UCSC Table browser (http://genome.ucsc.edu/). Proximal promoters were defined as ±10kb from the transcription start site (TSS) and genic regions were identified as +10kbTSS to the transcription end site (TES). Peaks were assigned to genes if they overlapped the promoter or genic region by a minimum of 1bp using intersectBed. Gene body heatmaps and average density plots were generated using ngs.plot ^70^.

### Gene set enrichment analysis

Gene Set Enrichment Analysis (GSEA) was performed using GSEA/MSigDB^69^ HALLMARK and KEGG pathways based on ensemble gene lists from peaks within −10kb-TES for bivalent domains and ±10kbTSS for broad H3K4me3 domains. All pathways are significant based on FDR q-value.

### RNA-sequencing data processing

For TCGA data, raw counts were obtained from TCGAbiolinks^63^ in R (http://www.rstudio.com/).and transformed to TPM. For Roadmap data, raw counts were obtained from http://www.roadmapepigenomics.org/ and transformed to TPM. For all RNA-seq boxplots, gene expression values from both datasets were normalized using quantile normalization function in R. For identification of differentially expressed genes and GSEA analysis between NRAS and BRAF melanoma tumors, raw counts were obtained from TCGAbiolinks. DESeq2^71^ was employed for normalization of differential gene expression analysis. GSEA^69^ was run on a ranked gene list using HALLMARK gene sets with default settings.

### Data and software availability

The ChIP and mRNA sequencing data have been deposited in the NCBI GEO BioProject database with the following accession number GSE134043.

