## Supplemental_Information for "Bivalent and Broad Chromatin Domains Regulate Pro-metastatic Drivers in Melanoma"

### Figure S1

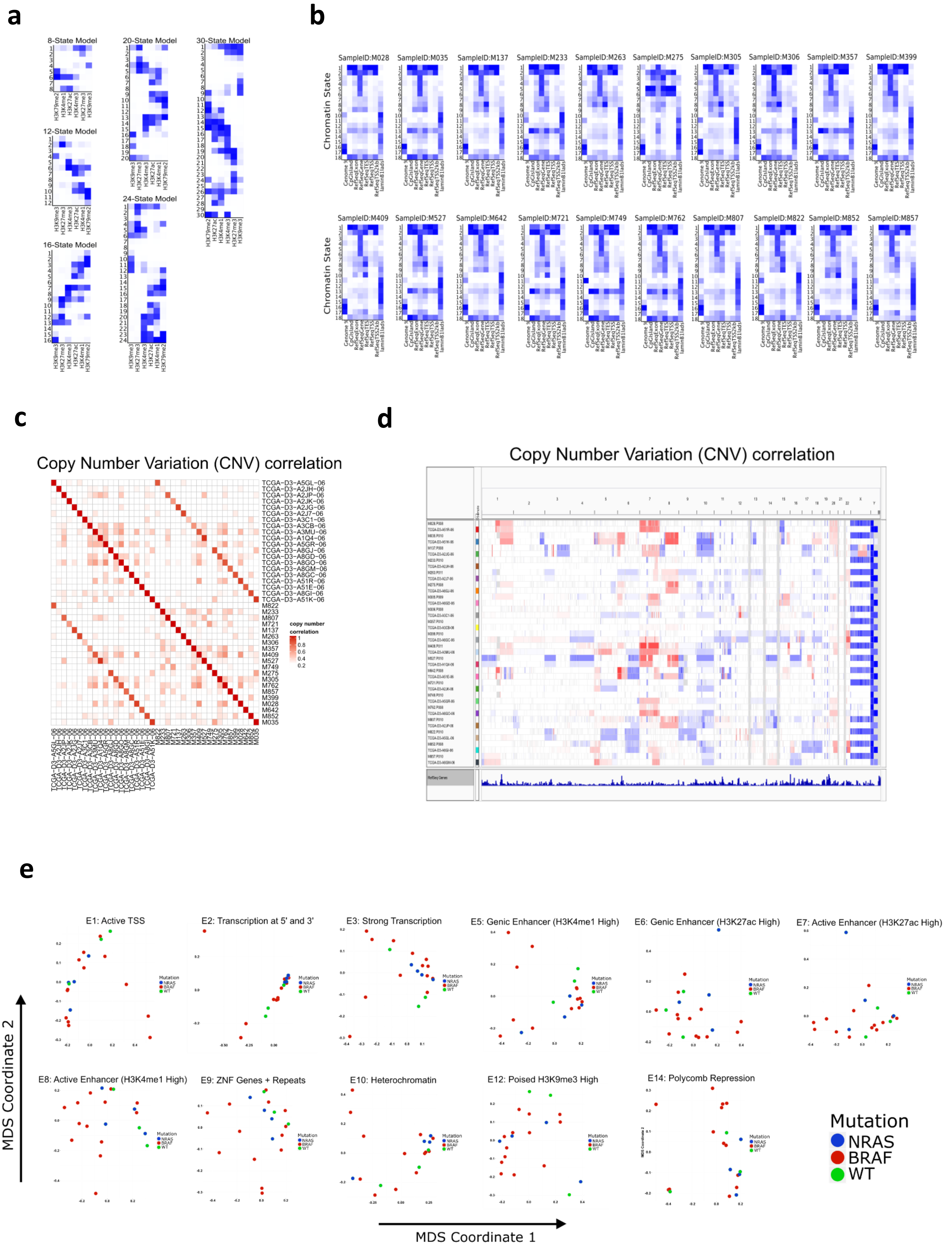


**Figure S1:**
a) Representative combinatorial chromatin state definitions and histone mark probabilities identified in 20 metastatic melanoma tumor samples using the ChromHMM algorithm.
b) Genomic annotation enrichments for each chromatin state in 20 metastatic melanoma tumor samples.
c) Copy Number Variation (CNV) correlation between the 20 melanoma tumors profiled and CNV classification from TCGA based on all genes.
d) Genome browser view of CNV correlation from (c).
e) MDS analysis of chromatin states annotated by mutation (NRAS, BRAF and WT) classifications from TCGA.

### Figure S2

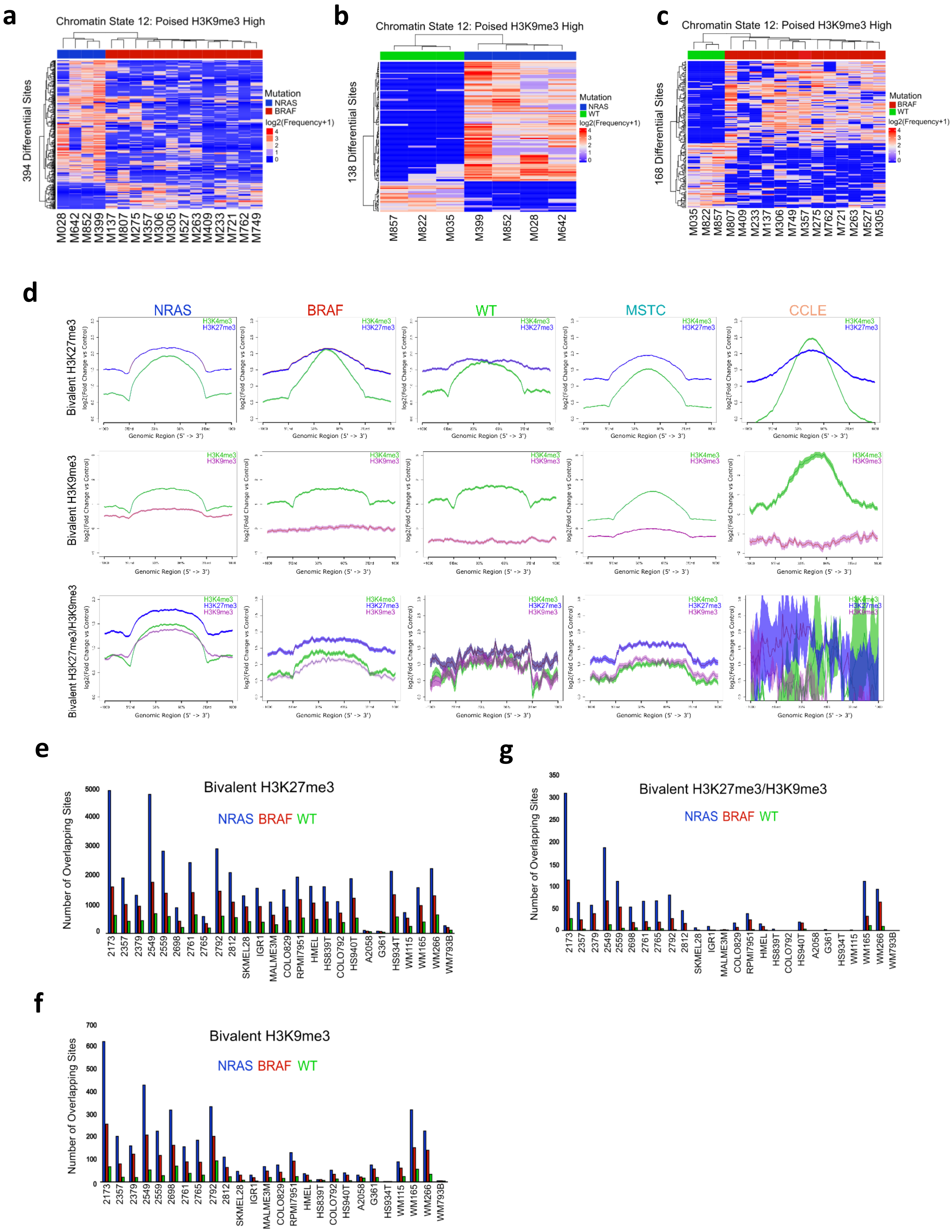


**Figure S2:**

a-c) Heatmaps displaying differentially regulated regions (FDR < 0.05) of chromatin state 12 (poised H3K9me3 high) between NRAS, BRAF and WT tumor subtypes.

d) Average density profiles of common bivalent H3K27me3, bivalent H3K9me3 and bivalent H3K27/H3K9me3 loci in melanoma tumor subtypes, MSTC and CCLE lines.

e) Co-occupancy analysis of common bivalent H3K27me3 binding sites in melanoma tumor subtypes directly overlapping bivalent H3K27me3 binding sites in individual MSTC or CCLE lines.

f) Co-occupancy analysis of common bivalent H3K9me3 binding sites in melanoma tumor subtypes directly overlapping bivalent H3K9me3 binding sites in individual MSTC or CCLE lines.

g) Co-occupancy analysis of common bivalent H3K27/H3K9me3 binding sites in melanoma tumor subtypes directly overlapping total bivalent H3K27/H3K9me3 binding sites in individual MSTC or CCLE lines.

### Figure S3

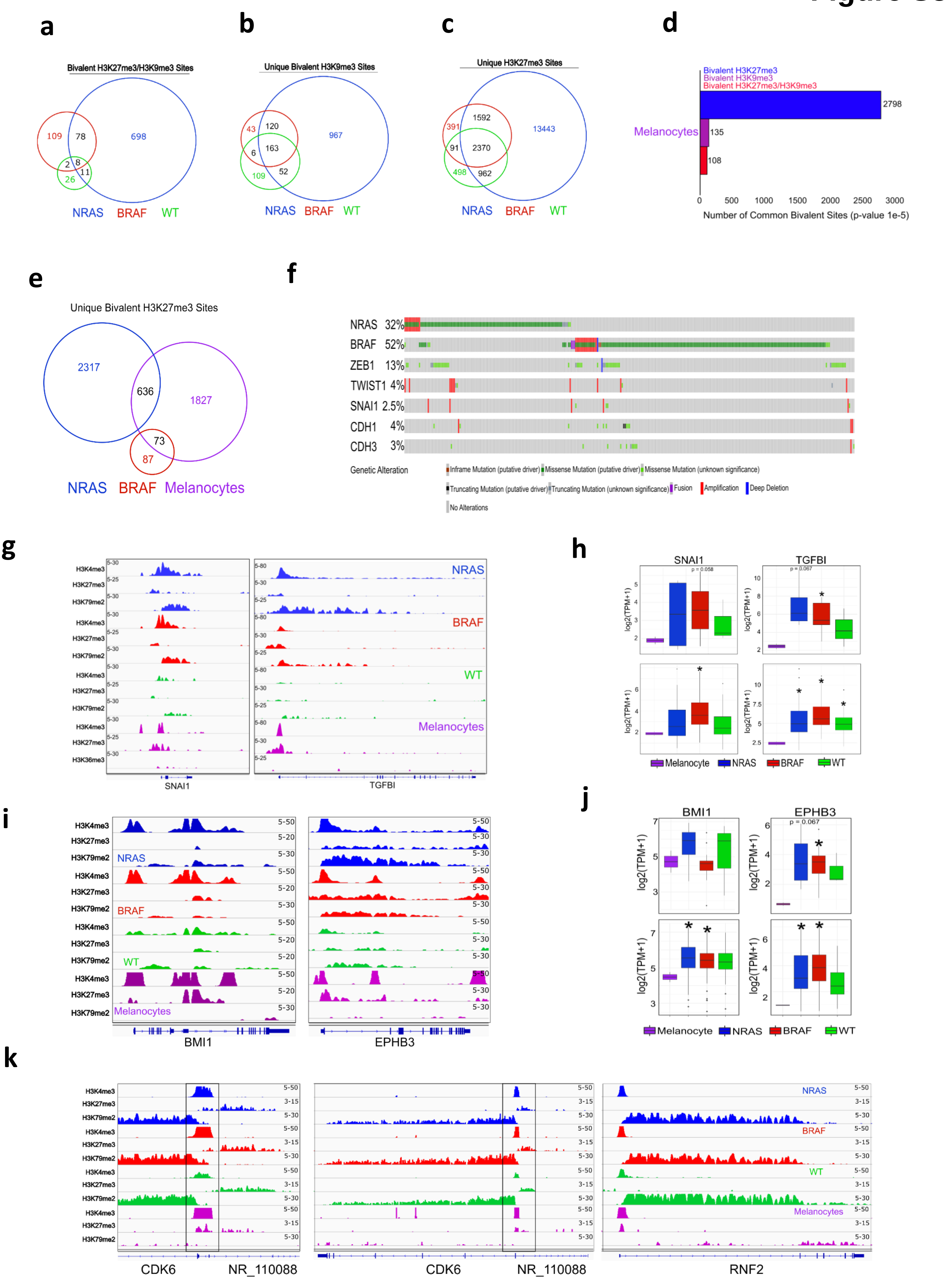

**Figure S3:**

a-c) Venn diagram analysis of bivalent H3K27/H3K9me3, unique bivalent H3K9me3 and unique H3K27me3 binding sites in melanoma tumor subtypes.

d) Common bivalent H3K27me3, bivalent H3K9me3 and bivalent H3K27/H3K9me3 binding sites in melanocytes. Binding sites were identified as common if they were present in 100% of the samples (2/2 melanocytes).

e) Venn diagram analysis of shared bivalent H3K27me3 sites in NRAS and BRAF tumor subtypes overlapping bivalent H3K27me3 sites in primary melanocytes from Roadmap.

f) OncoPrint of all possible genetic alterations in NRAS, BRAF, ZEB1, TWIST1, SNAI1, CDH1 and CHD3 genes from 363 skin cutaneous melanoma samples.

g) Genome browser view of ChIP-seq tracks for H3K4me3, H3K27me3 and active transcription (H3K79me2/H3K36me3) on the *SNAI1 and TGFBI* genes in melanocytes and melanoma tumor subtypes.

h) Boxplot displaying quantile normalized mean RNA-expression profiles (log2 TPM) of the *SNAI1 and TGFBI* genes in melanocytes (n=2) and melanoma tumor subtypes (NRAS=4, BRAF=13, WT=3 (top), NRAS=81, BRAF=118, WT=38 (bottom). Asterisk denotes p-value < 0.05.

i) Genome browser view of ChIP-seq tracks for H3K4me3, H3K27me3 and active transcription (H3K79me2/H3K36me3) on the *BMI1 and EPHB3* genes in melanocytes and melanoma tumor subtypes.

j) Boxplot displaying quantile normalized mean RNA-expression profiles (log2 TPM) of the *BMI1 and EPHB3* genes in melanocytes (n=2) and melanoma tumor subtypes (NRAS=4, BRAF=13, WT=3 (top), NRAS=81, BRAF=118, WT=38 (bottom). Asterisk denotes p-value < 0.05.

k) Genome browser view of ChIP-seq tracks for H3K4me3, H3K27me3 and active transcription (H3K79me2/H3K36me3) on the *CDK6 and RNF2* genes in melanocytes and melanoma tumor subtypes.

#### Figure S4

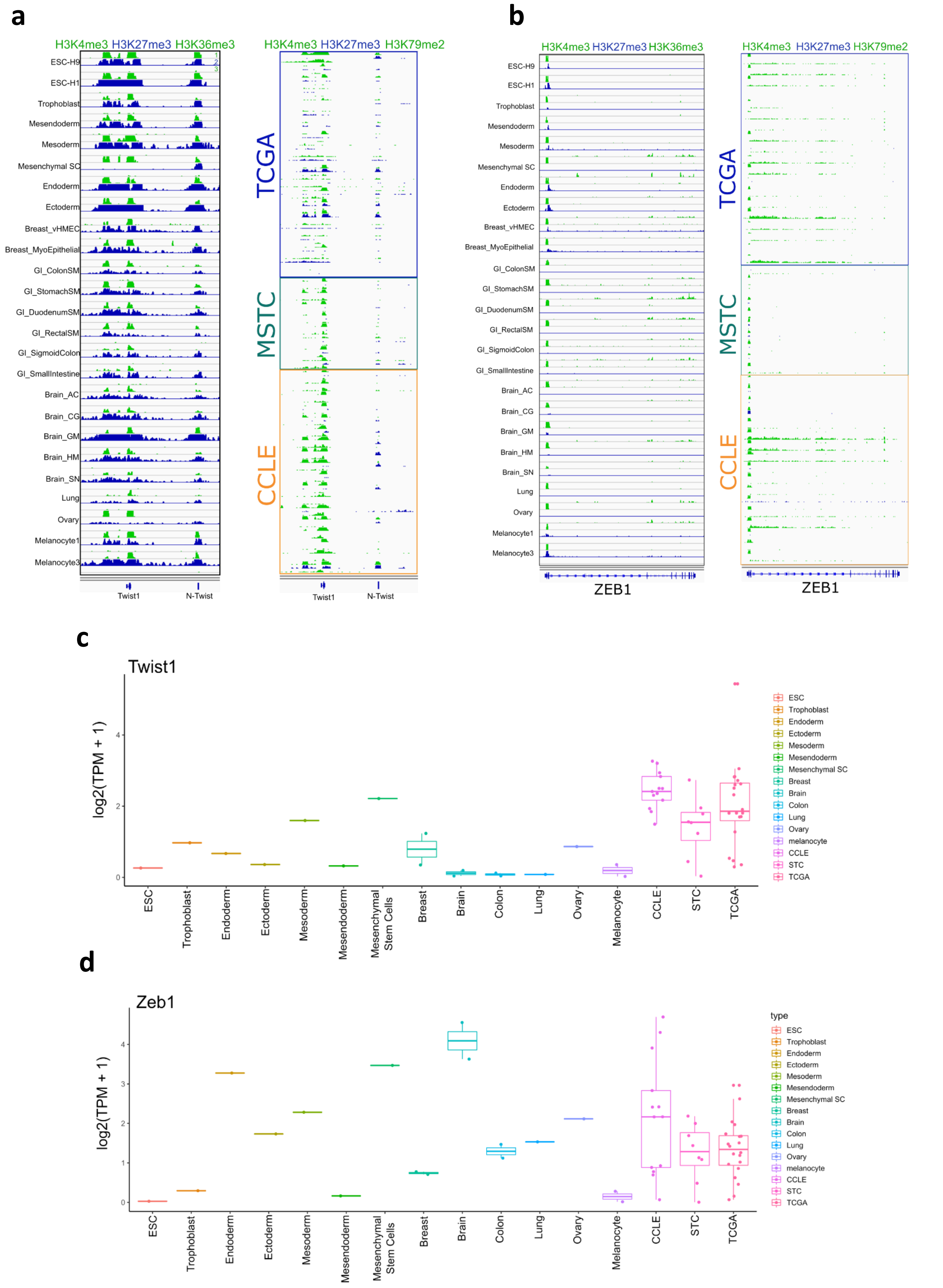


**Figure S4:**
a, b) Genome browser view of ChIP-seq tracks for H3K4me3, H3K27me3 and active transcription (H3K79me2/H3K36me3) on the *TWIST1* and *ZEB1* genes in ESC, Trophoblast, Ectoderm, Endoderm, Mesoderm, Mesendoderm, Mesenchymal Stem Cells, normal Breast, Brain, Colon, Lung, Ovary and melanocytes tissues or cell lines, as well as melanoma tumors, MSTC and CCLE.
c, d) Boxplots displaying quantile normalized mean RNA-expression profiles (log2 TPM) of the *TWIST1 and ZEB1* genes in ESC, Trophoblast, Ectoderm, Endoderm, Mesoderm, Mesendoderm, Mesenchymal Stem Cells, normal Breast, Brain, Colon, Lung, Ovary and melanocytes tissues or cell lines, as well as melanoma tumors, MSTC and CCLE.

Figure S5

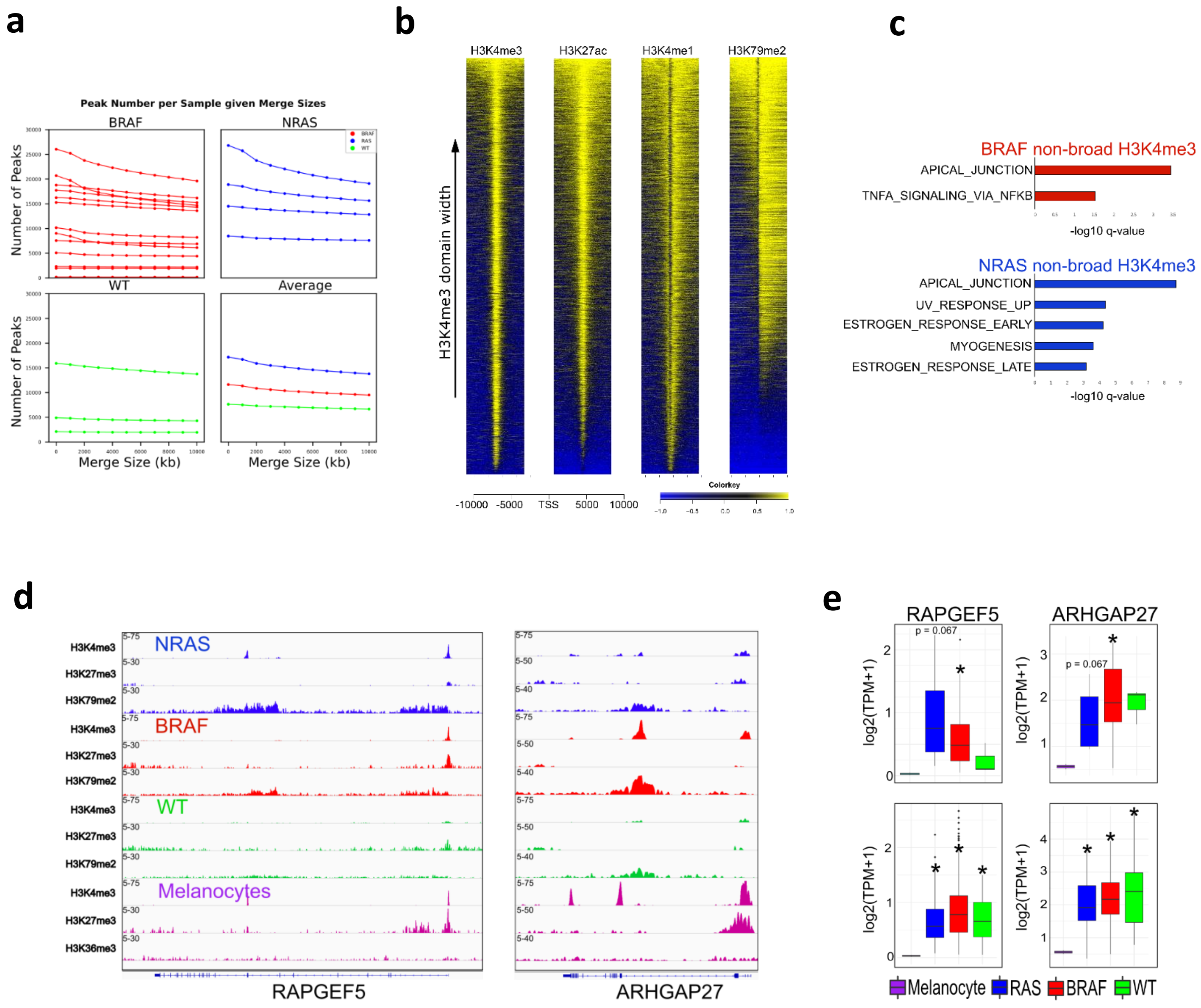

**Figure S5:**
a) LinePlot displaying total H3K4me3 peak number at merge sizes between 1kb to 10kb in BRAF, NRAS and WT samples. Bottom right plot displays average by mutational subtype.
b) Average density profile displaying broad (>4kb) and non-broad (<4kb) H3K4me3, H3K27ac, H3K4me1 and H3K79me3 domains at -10kb to +10kb around TSS. Heatmap sites for all marks were based on H3K4me3 associated genes.
c) Top significant MSigDB/GSEA HALLMARK pathways based on bivalent H3K27me3 genes (-/+10kbTSS-TES) that are lost in melanocytes and associated with non-broad H3K4me3 in NRAS, BRAF and WT tumor subtypes identified in Fig.5a.
d) Genome browser view of ChIP-seq tracks for H3K4me3, H3K27me3 and active transcription (H3K79me2/H3K36me3) on the *RAPGEF5* and *ARHGAP27* genes in melanocytes and melanoma tumors.
e) Boxplot displaying quantile normalized mean RNA-expression profiles (log2 TPM) of the *RAPGEF5* and *ARHGAP27* genes in melanocytes (n=2) and melanoma tumor subtypes (NRAS=4, BRAF=13, WT=3 (top), NRAS=81, BRAF=118, WT=38 (bottom). Asterisk denotes p-value < 0.05.

### Figure S6

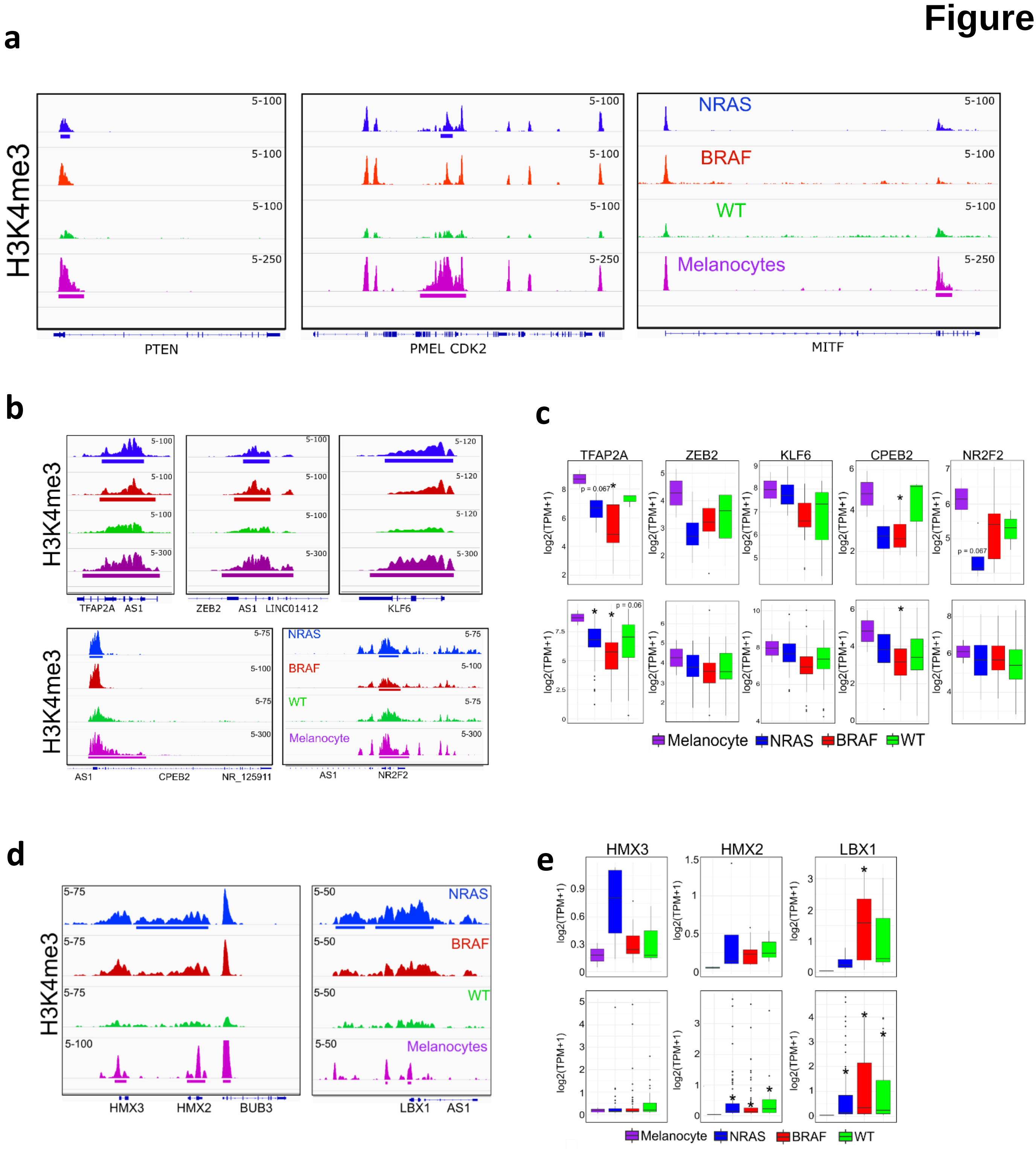


**Figure S6:**

a) Expanded genome browser view of ChIP-seq tracks displaying H3K4me3 shortening on the *PTEN*, *PMEL* and *MITF* genes in melanoma tumors relative to melanocytes.

b) Genome browser views of ChIP-seq tracks displaying H3K4me3 shortening on the *TFAP2A*, *ZEB2*, *KLF6*, *CPEB2* and *NR2F3* genes in melanoma tumors relative to melanocytes.

c) Boxplot displaying quantile normalized mean RNA-expression profiles (log2 TPM) of the *TFAP2A*, *ZEB2*, *KLF6*, *CPEB2* and *NR2F3* genes in melanocytes (n=2) and melanoma tumor subtypes (NRAS=4, BRAF=13, WT=3 (top), NRAS=81, BRAF=118, WT=38 (bottom).

d) Genome browser views of ChIP-seq tracks displaying H3K4me3 lengthening on the *HMX2/3* and *LBX1* genes in melanoma tumors relative to melanocytes.

e) Boxplot displaying quantile normalized mean RNA-expression profiles (log2 TPM) of the *HMX2/3* and *LBX1* genes in melanocytes (n=2) and melanoma tumor subtypes (NRAS=4, BRAF=13, WT=3 (top), NRAS=81, BRAF=118, WT=38 (bottom). Asterisk denotes p-value < 0.05.

### Figure S7

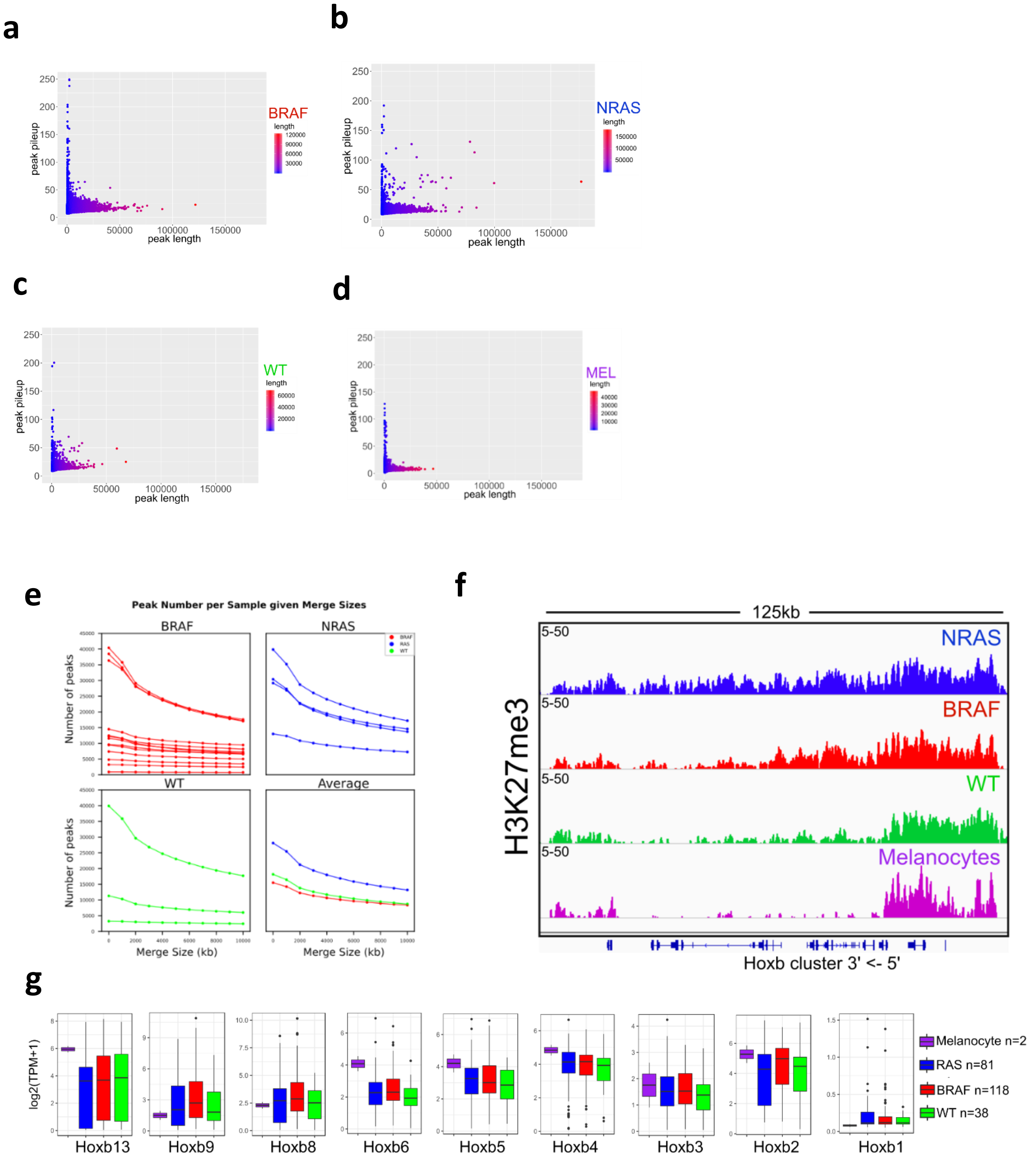


**Figure S7:**
a) Scatterplots of peak width (x-axis) and height (y-axis) from MACS2 broad peak calls (p-value 1e-5) for H3K27me3 in BRAF, (b) NRAS, and (c) WT samples.
d) Scatterplots of peak width (x-axis) and height (y-axis) from MACS2 broad peak calls (p-value 1e-5) in melanocytes.
e) LinePlot displaying total H3K27me3 peak number at merge sizes between 1kb to 10kb in BRAF, NRAS and WT samples. Bottom right plot displays average by mutational subtype.
f) Genome browser views of ChIP-seq tracks displaying exceptionally wide H3K27me3 lengthening from the 5’->3’ end on *HOXB* cluster genes in BRAF mutational subtypes.
g) Boxplot displaying quantile normalized mean RNA-expression profiles (log2 TPM) of all *HOXB* cluster genes in melanocytes (n=2) and melanoma tumor subtypes (NRAS=81, BRAF=118, WT=38). Asterisk denotes p-value < 0.05.
